# Lysophosphatidyl-choline 16:0 mediates persistent joint pain through Acid-Sensing Ion Channel 3: preclinical and clinical evidences

**DOI:** 10.1101/2021.03.29.437487

**Authors:** Florian Jacquot, Spiro Khoury, Bonnie Labrum, Kévin Delanoe, Ludivine Pidoux, Julie Barbier, Lauriane Delay, Agathe Bayle, Youssef Aissouni, David A. Barriere, Kim Kultima, Eva Freyhult, Anders Hugo, Eva Kosek, Aisha S. Ahmed, Alexandra Jurczak, Eric Lingueglia, Camilla I. Svensson, Véronique Breuil, Thierry Ferreira, Fabien Marchand, Emmanuel Deval

## Abstract

Rheumatic diseases are often associated to debilitating chronic joint pain, which remains difficult to treat and requires new therapeutic strategies. Here, we describe increased content of lysophosphatidyl-choline (LPC) 16:0 in the knee synovial fluids of two independent cohorts of patients with painful joint diseases. If LPC16:0 levels correlated with pain in patients with osteoarthritis (OA), they do not appear to be the hallmark of a particular joint disease. We found that intra-articular injections of LPC16:0 in mouse produce chronic pain and anxiety-like behaviors in both males and females with no apparent inflammation, peripheral nerve sprouting and damage, nor bone alterations. LPC16:0-induced persistent pain state is dependent on peripheral Acid-Sensing Ion Channel 3 (ASIC3), ultimately leading to central sensitization. LPC16:0 and ASIC3 thus appear as key players of chronic joint pain with potential implications in OA and possibly across others rheumatic diseases.

## Introduction

Chronic joint pain is the primary reason why patients seek care from rheumatologists, representing a heavy burden for modern societies^1,2^. In addition to economical costs, chronic joint pain considerably impairs the patients’ quality of life, leading to disabilities and psychological distress^1,3^. Despite major advances in inflammatory arthritis therapies, most patients, especially those suffering from osteoarthritis (OA), continue to experience life-disabling pain. The development of new therapeutic strategies requires a better understanding of the pathophysiology of joint pain, with careful identification of the triggering factors and associated mechanisms.

In the last two decades, signaling lipids have been shown to contribute to the onset and maintenance of pain. Lysophosphatidyl-choline (LPC) and lysophosphatidic acid (LPA), the most prominent lyso-glycero-phospholipids^4^, are emerging as potential direct pain mediators, besides well-known thromboxanes, leukotrienes and prostaglandins^5,6^. LPA has been already identified as a key promoter of nerve demyelination and certainly of neuropathic pain^7,8^. Paradoxically, LPA receptor signaling has also documented physiological roles in myelinating cells during nervous system development^9–11^. The role of LPC in pain remains more elusive, especially *in vivo*, since its effects could be partially mediated by LPA through the action of autotaxin, the LPC-to-LPA conversion enzyme^12,13^. Nevertheless, some studies support a direct role of LPC in pain associated to its ability to activate/potentiate pain-related ion channels^5,6,14–16^. Moreover, LPC could be particularly important in pain originating from deep tissues such as joints^6,17^ and muscles^18^. Concerning joints, we already reported high levels of LPC in human synovial fluids from patients with painful joint diseases^6^ and more recently, an over activation of the conversion pathway of PC-to-LPC has been observed in patients with OA^17^.

LPC is able to activate and potentiate pain-related Acid-Sensing Ion Channels 3 (ASIC3)^6^, a sub-type of the ASIC channel family, widely expressed in nociceptors and also gated by extracellular protons^19^. ASIC3 have been involved in animal models of inflammatory and non-inflammatory pain^20,21^, especially in the development of chronic pain states originating from joints^22–24^ and muscles^18,25,26^. Accordingly, ASIC3-dependent pain behaviors can be induced in rodent upon injection of LPC in the paw or the muscle^6,18^. Here we investigated the clinical relevance of LPC in chronic joint pain and we explored its mechanism of action that involved peripheral ASIC3 channels, using a new preclinical pain model in mice induced by intra-articular administrations of LPC 16:0.

## Results

### Patients with painful joint diseases exhibit higher levels of knee synovial LPC16:0

The knee synovial fluids of 35 patients from a first cohort (OA patients only, **supplementary table 1**) displayed significantly higher concentrations of LPC compared to controls (77.4±5.0 μM and 40.0±3.4 μM, respectively; **fig.1a**). The distribution of fatty acyl chains within this lipid class revealed that LPC16:0, bearing the long palmitate saturated chain, was the most represented species in both control and OA individuals (**fig. 1b**). Moreover, LPC16:0 was the only species significantly increased in OA samples compared to controls (35.2±2.4 μM *vs* 16.3±1.5 μM; **fig. 1b**), indicating that the higher contents of LPC found in patients suffering from OA were mainly supported by an increase of LPC16:0 in their synovial fluids.

**Figure 1:**
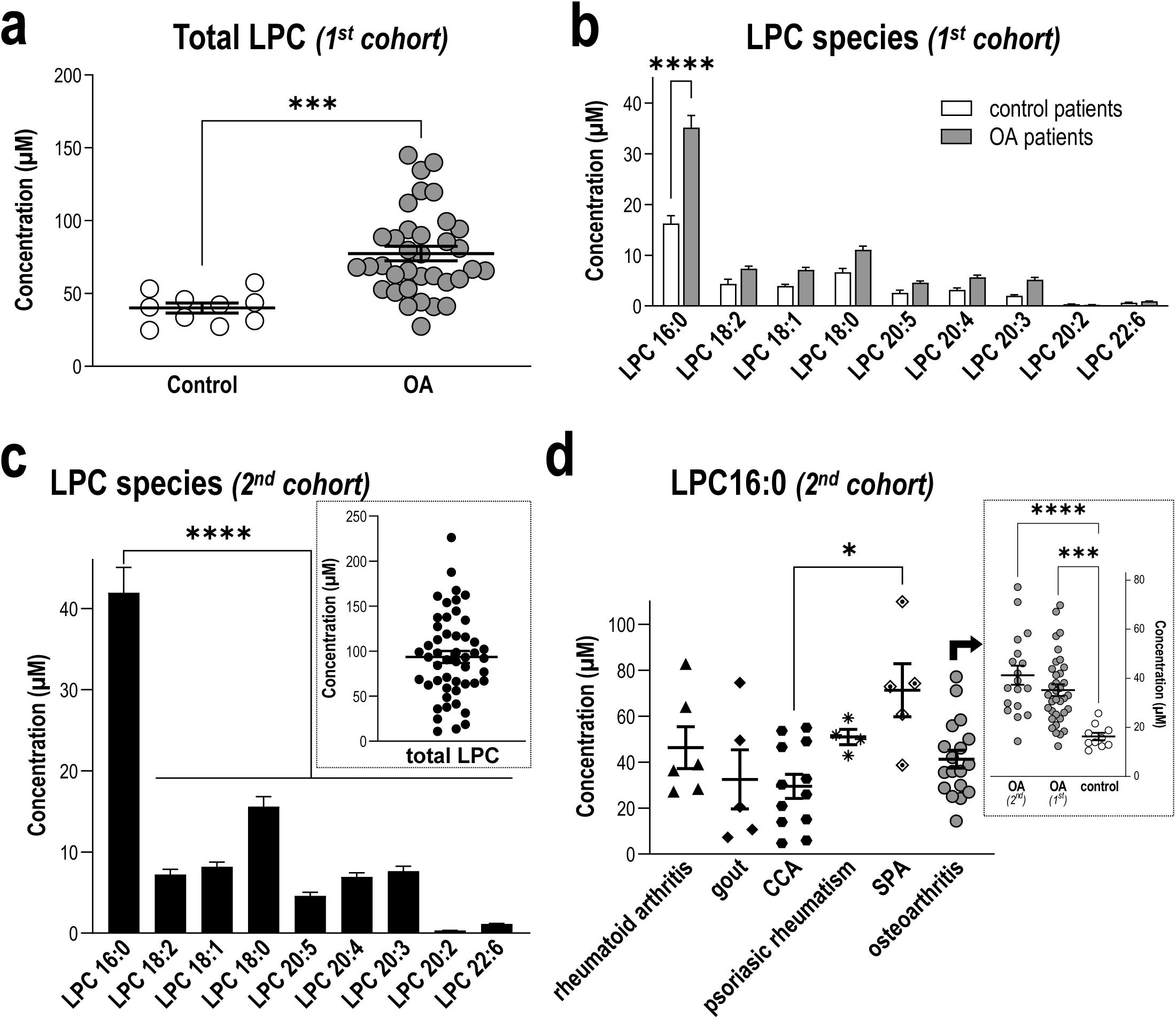
Levels of LPC16:0 are increased in synovial fluids of patients suffering from joint pain. Quantifications of LPC species were performed in knee human synovial fluid (HSF) samples. Total lipids were extracted from HSF samples and analyzed by direct infusion in mass spectrometry (MS) using an electrospray ionization source (ESI) in the positive ion mode (see “Methods” section for details). **a**, Comparison of total LPC concentrations (μM) between a first cohort of patients with osteoarthritis (OA, n=35 patients) and post mortem controls (n=10), showing significant higher levels of LPC in patients compared to controls (\*\*\**p*=0,0003, Unpaired *t* test). **b**, Distribution of the different LPC species concentrations (μM) in the HSF of OA patients and post mortem controls. Although mean concentrations of most LPC species were higher in the HSF of OA patients, LPC16:0 was the most abundant and the only one to be significantly elevated compared to post mortem controls (\*\*\**p*<0.0001, two-way ANOVA followed by a Sidak’s multiple comparison test). **c**, Distribution of the different LPC species concentrations (μM) in the HSF of a second cohort of patients suffering from different joint pathologies (n=50), showing significant higher level of LPC16:0 compared to the others species (\*\*\*\**p*<0.0001, One-way ANOVA followed by a Dunnet’s multiple comparison test). *Inset:* Total LPC concentration in HSF of the second cohort of patients. **d**, LPC16:0 concentrations in the different subgroups of patients from the second cohort: rheumatoid arthritis (RA, n=6), gout (n=5), chondrocalcinosis (CCA, n=12), psoriatic arthritis (PA, n=4), spondyloarthritis (SPA, n=5), and OA (n=18). The LPC16:0 concentrations in all the different subgroups of patients are comparable, except a difference between CCA and SPA (*, *p*<0.05, Kruskal-Wallis test followed by a Dunn’s multiple comparison test). *Inset:* LPC16:0 concentrations in HSF of OA patients from the first and second cohort are similar and higher than control post mortem (\*\*\**p*<0.001 and \*\*\*\**p*<0.0001, One-way ANOVA followed by a Tukey’s multiple comparison test).

Correlation studies were carried out between LPC16:0 concentrations and pain outcomes, including Visual Analog Scores (global and knee VAS) and Knee injury Osteoarthritis Outcome Score (KOOS), with data adjusted for patients’ age, gender, body mass index (BMI) and synovial fluid interleukin 6 (IL-6) level (**supplementary table 2**). Significant correlations between LPC16:0 concentrations and pain (VAS) were observed (*p*=0.016 or *p*=0.031 for VAS knee and VAS global, respectively, **supplementary table 2**), and an inverse correlation with KOOS questionnaire almost reached statistical significance (*p*=0.060, **supplementary table 2**). Including or excluding IL-6 levels in the ANOVA did not have drastic impact on the association between LPC16:0 and pain measures (*p*=0.016/0.019, *p*=0.031/0.066 and *p*=0.060/0.080 for VAS knee, VAS global and KOOS, respectively, including/excluding IL-6). In line with this, there were no significant correlations between LPC16:0 and IL-6 levels in synovial fluids (**supplementary table 2)**, indicating that the involvement of LPC16:0 in patient pain outcomes (at least for VAS knee and VAS global) was not associated to an IL6-dependent inflammation in this first cohort of OA patients.

To confirm and expand these results, we next assessed whether increase of LPC16:0 knee synovial fluids’ levels could be the hallmark of joint pathologies associated or not with *bona fide* inflammation by measuring LPC concentrations in the synovial fluids from a second heterogeneous cohort of patients (**supplementary table 3**) with various inflammatory and non-inflammatory joint diseases (**fig. 1c–d**). The data pooled from 50 patients, irrespective of the disease, showed a distribution of LPC species (**fig. 1c**) that was similar to the first cohort of patients with OA (**fig 1b**), with an average concentration for total LPC of 93.6±6.7 μM (**fig. 1c, inset**), significantly higher than controls samples (**fig. 1a**, *p*=0.0007, Unpaired *t* test test). LPC16:0 was also the major contributor to this elevated LPC level, as for the first OA cohort, with an average concentration of 41.9±3.1 μM (**fig. 1c)**, which was significantly higher than control samples (**fig. 1b**, *p*=0.0006, Unpaired *t* test). LPC16:0 concentrations were then pooled, according to the patients’ diseases as follow (**fig. 1d**): rheumatoid arthritis (RA, n=6), gout (n=5), chondrocalcinosis (CCA, n=12), psoriatic arthritis (PA, n=4), spondyloarthritis (SPA, n=5) and OA (n=18). LPC16:0 levels in these different subgroups of patients were similar, except a difference between CCA and SPA patients. Although, LPC16:0 did not appear to be the hallmark of a particular joint pathology, patients with micro-crystalline arthropathies (CCA and gout) displayed lower levels of synovial LPC16:0 (**supplementary fig. 1**) than patients suffering from other inflammatory rheumatic diseases (RA + SPA + PA), suggesting that LPC16:0 might have a less prominent role in pain associated with these two particular joint pathologies. Most importantly, OA patients from this second cohort also displayed an elevated level of LPC16:0 that was similar to the first OA cohort and significantly different from control patients (**fig. 1d, inset**).

### Intra-articular LPC16:0 injections induce persistent pain and anxiety-like behaviors in mice

As LPC16:0 concentration was higher in the synovial fluids of patients suffering from painful joint diseases compared to controls (**fig. 1**), we investigated the potential pronociceptive effect, as well as associated comorbidities such as anxiety-like behaviors, following ankle intra-articular injections of LPC16:0 in mice (**fig. 2a–e and supplementary fig. 2**). Comparatively, a single complete freund adjuvant (CFA) injection into the ankle joint was used as a positive control. As expected, intra-articular injection of CFA induced a significant decrease of mechanical withdrawal threshold only in the ipsilateral paw (**fig. 2b and supplementary fig. 2a**) from day 1 through day 28 after CFA injection, with the maximal intensity at day 14. CFA-induced mechanical allodynia was also associated with a significant weight bearing deficit (**fig. 2c**). LPC16:0 was injected twice intra-articularly on day 0 and day 5 into the ankle joint (**figs. 2a–e and supplementary fig 2**), at the dose of 10 nmol, which corresponded to the amount of lipids we had previously shown to generate acute pain-like behaviors when injected at the plantar surface in mice^6^, and was in the lower range of LPC16:0 quantities assessed here in patient synovial fluids. The first injection induced a significant short-lasting mechanical allodynia (up to day 5), only on the ipsilateral side compared to vehicle (**fig. 2b and supplementary fig 2a**). The second injection of LPC16:0 led to a significant long-lasting ipsilateral mechanical allodynia until the end of the protocol on day 28. This persistent allodynia was similar to the effect of CFA, and was also associated with altered weight-bearing up to day 21 (**fig. 2c**). A significant thermal hyperalgesia was also observed in LPC16:0-injected mice, with a sustained effect similar to CFA (**supplementary fig. 2b)**. In contrast, intra-articular injections of vehicle did not change mechanical or heat paw withdrawal thresholds (**fig. 2b and supplementary fig. 2a**), nor distribution of weight bearing over the animal’s two hindpaws (**fig. 2c**). Joint pain is often associated with comorbidities such as stress and anxiety^3^, and we therefore assessed the development of anxiety-like behaviors following intra-articular injections of LPC16:0 in mice. In the open field test, administration of LPC16:0, as well as CFA, did not alter locomotor behavior (distance traveled) compared to control mice (**supplementary fig. 2c**), but induced a significant decrease in the time spent in the center (**fig. 2d**), indicating the development of increased thigmotactism. A decrease of the time spent in the open arm in the elevated plus maze test was also observed in both LPC16:0 and CFA groups compared to control animals (**fig. 2e)**, demonstrating the development of anxiety-like behaviors. This result was further reinforced by the hole board and marble burying tests, where a decrease in the number of head dips and an increase of buried marble were observed, respectively, following LPC16:0 and CFA administrations (**supplementary fig. 2d-e**). The development of anxiety-like behaviors was associated to increase neuronal activity within the amygdala, particularly in the basolateral nucleus (BLA), in LPC-treated mice compared to controls, as shown by c-fos immunochemistry (**supplementary fig. 3a-c**). Importantly, the effect of LPC16:0 was not due to its conversion to LPA by autotaxin, since neither co-nor post-treatment with an autotaxin inhibitor (S32826, intra-articular injection) altered the pronociceptive effect of LPC16:0 intra-articular injections (**fig. 2f–g)**.

**Figure 2:**
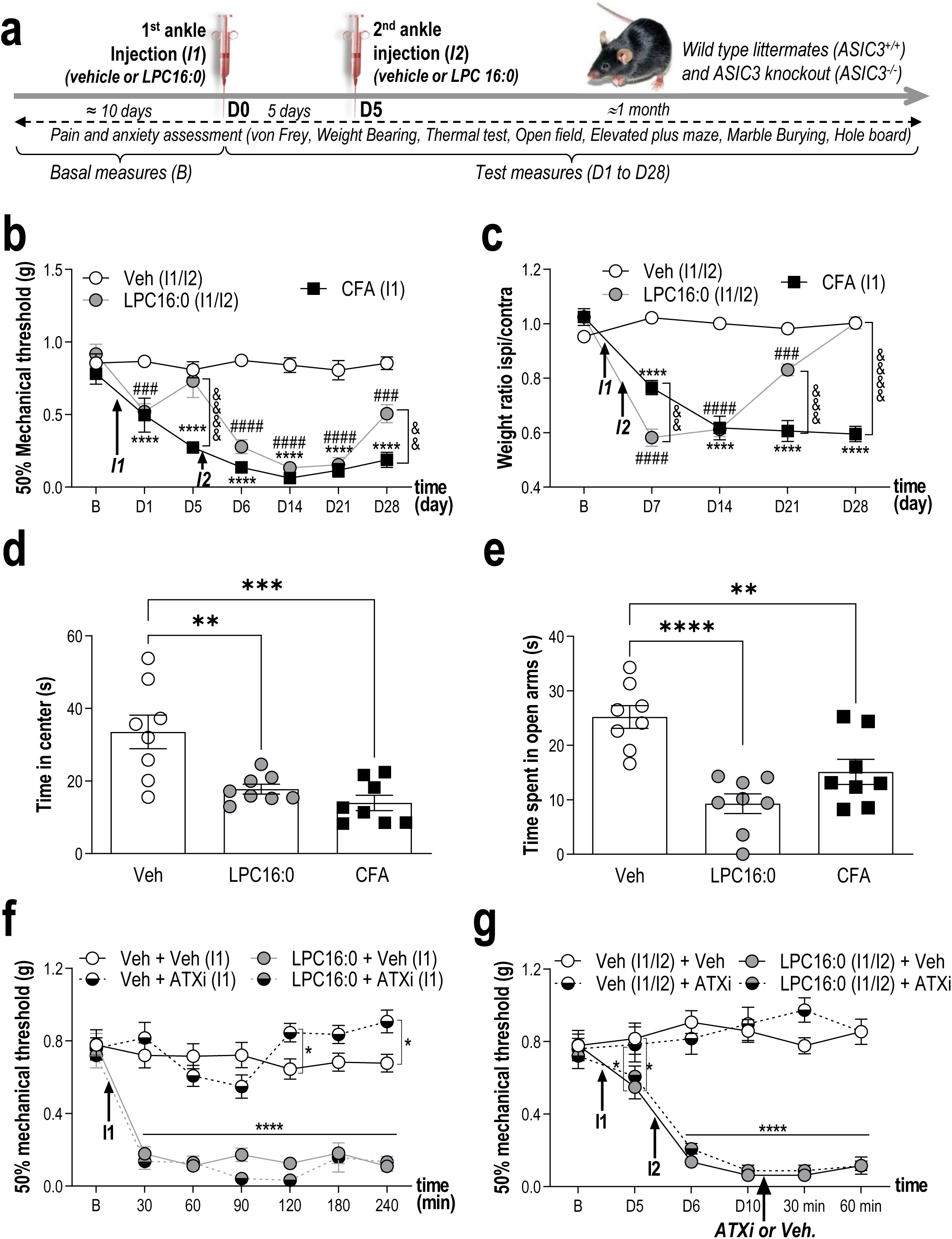
LPC injections into mouse ankle joints produce long-lasting pain-like behaviors associated with anxiety-related behaviors. **a**, Timeline of the experimental procedures used. **b**, Effect of intra-articular ankle injections of LPC16:0 (10 nmol), complete freund adjuvant (CFA) or vehicle (Veh) on mechanical allodynia in male mice. Ipsilateral mechanical paw withdrawal threshold was assessed using the up and down method with von Frey filament from D1 to D28. Results are expressed as 50% mechanical threshold (n=8-16 mice per group; \*\*\*\**p*<0.0001 for Veh *vs.* CFA; ###*p*<0.001 and ^####^*p*<0.0001 for Veh *vs.* LPC16:0; ^&&^*p*<0.01 and ^&&&&^*p*<0.0001 for CFA *vs.* LPC16:0; Two-way ANOVA followed by a Tukey’s *post hoc* test). **c**, Effect of intra-articular injections of LPC16:0 (10 nmol), CFA or vehicle on weight bearing in male mice. Results are expressed as the weight ratio between the ipsilateral and contralateral hindpaws (n=8 mice per group; \*\*\*\**p*<0.0001 for Veh *vs.* CFA; ^###^*p*<0.001 and ^####^*p*<0.0001 for Veh *vs.* LPC16:0; ^&&^*p*<0.01, ^&&&^*p*<0.001 and ^&&&&^*p*<0.0001 for CFA *vs*. LPC16:0; Two-way ANOVA followed by a Tukey’s *post hoc* test). **d-e**, Effect of intra-articular injections of LPC16:0 (10 nmol), CFA or vehicle on the time spent in the center of an open field test (**d**) and on the time spent in the open arms in the elevated plus maze test (**e**). One-way ANOVA followed by Tukey’s *post hoc* tests with \*\**p*<0.01, \*\*\**p*<0.001 and \*\*\**p*<0.0001 (n=8 mice per group). **f**, The autotaxin inhibitor S32826 (ATXi) was co-injected at the dose of 10 nmol with LPC16:0 (10 nmol, I1) and mechanical paw withdrawal thresholds were assessed from 30 min to 240 min post-injection (n=8 mice per group, \**p*<0.05 and ****p<0,0001, two-way ANOVA followed by a Tukey’s *post hoc* test). **g**, S32826 was injected at D10, *i.e.*, after the two injections of LPC16:0 (10 nmol) 5 days apart (I1/I2), in the same ankle joint, and mechanical paw withdrawal thresholds were assessed at 30 and 60 min post-injection (n=8 per group, \**p*<0.05 and \*\*\*\**p*<0,0001, Two-way ANOVA followed by a Tukey’s *post hoc* test).

**Figure 3:**
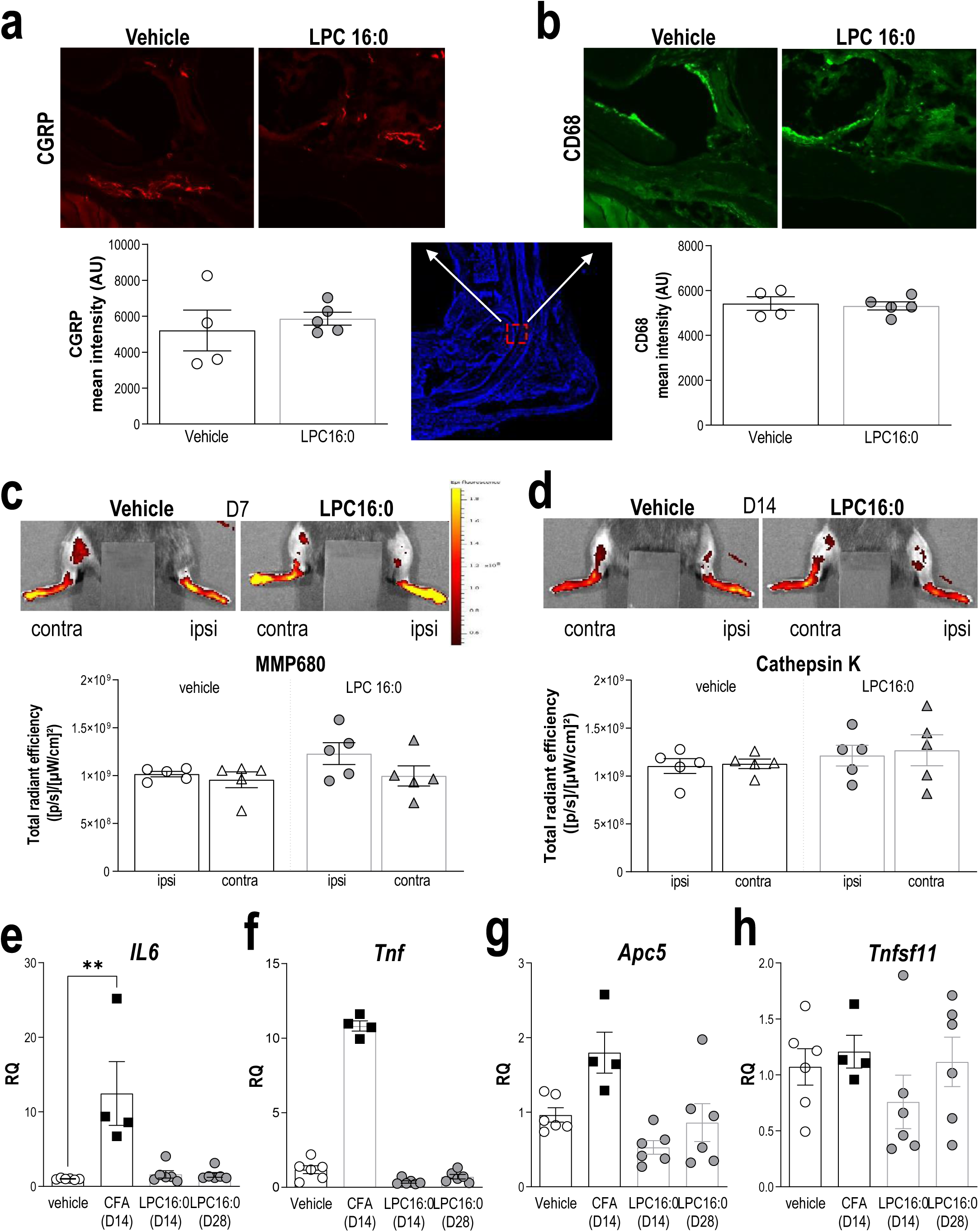
LPC16:0 injections into mouse ankle are not associated to peripheral nerve sprouting, inflammation, nor bone alterations. **a-b**, Analysis of CGRP and CD68 expression in the ipsilateral tibiotarsal joint at D28, following intra-articular administrations of LPC16:0 (10 nmol) or vehicle in male mice. *Top*, representative photomicrographs of CGRP (**a**) and CD68 (**b**) staining in vehicle and LPC16:0-treated mice. *Bottom*, quantification of CGRP positive fibers (**a**) in two ROIs (570 × 570 μm), and of CD68 positive cells (**b**) in one ROI (2500 × 2500 μm) from the ipsilateral tibiotarsal joint of vehicle and LPC16:0-injected mice. Results are expressed as a mean intensity (n= 4 for each group, no significant differences, Mann-Whitney tests). **c-d**, Evaluation of the effect of LPC16:0 (10 nmol) or vehicle on inflammation and bone remodeling using *in vivo* fluorescent imaging for metalloprotease (MMP) activity at D7 (**c**) and Cathepsin K activity at D14 (**d**), respectively, in the ipsilateral and contralateral hindpaws of male mice. *Upper panels*, illustrations; *Lower panels*, MMP ratio quantification (**c**) and bone remodeling ratio quantification (**d**). Data from n=5 animals per group (no significant differences, Kruskal Wallis tests followed by a Dunn’s multiple comparison tests). **e-h**, Relative levels of *IL6* (**e**), *Tnf* (**f**), *Apc5* (TRAP) (**g**) and *Tnfsf11* (RANK-L) (**h**) mRNA expressed in the ipsilateral joint, determined by q-PCR after intra-articular injections of LPC16:0 (10 nmol, D14 and D28), CFA (D14) and vehicle (D14/D28; n=4-6 for each group, \*\**p*<0.01, Kruskal-Wallis tests followed by Dunn’s multiple comparison tests).

### Intra-articular LPC16:0 does not cause peripheral nerve sprouting, inflammation, nor bone alterations

We next investigated whether LPC16:0 ankle injections could generate any CGRP-positive nerve sprouting and/or CD68-positive cell infiltration in the synovium of the ipsilateral tibiotarsal joint (**fig. 3a–b)**. No difference in CGRP and CD68 staining was observed at day 28 between vehicle- and LPC16:0-injected mice, suggesting no peptidergic nerve fibers sprouting nor CD68^+^ macrophage lineage cells influx, respectively. A possible LPC16:0-induced joint inflammation was also evaluated using *in vivo* imaging with MMP680, a marker of matrix metalloprotease activity, as well as *Il6* and *Tnf* genes expression by q-PCR (**fig. 3c, e and f**). At day 7 (*i.e.*, two days after the second administration of LPC16:0), no increase of MMP680 fluorescence was observed in the ipsilateral joint of LPC16:0-injected mice compared to the contralateral joint and to joints of the control group (**fig. 3c**, photomicrograph and histogram). *II6* and *Tnf* mRNA levels in the ipsilateral joints of LPC16:0-injected mice were similar to the control group at day 14 and day 28, while an increase was observed in CFA-injected mice at day 14, as expected (**fig. 3e,f**). We also investigated if LPC16:0 could induce bone remodeling using *in vivo* imaging with Cathepsin K680, a marker of osteoclast activity, as well as the *Apc5* and *Tnfsf11* (coding for TRAP and RANK-L proteins, respectively) gene expression by q-PCR (**fig. 3d, g and h**). At day 14, Cathepsin K680 fluorescence was not increased in the ipsilateral joints of LPC16:0-injected mice compared to the contralateral ones and to joints of the control group (**fig. 3d**, photomicrograph and histogram). *Apc5* and *Tnfsf11* mRNA levels in the joints of LPC16:0 injected mice were similar to control mice at day 14 and day 28 (**fig. 3g,h**). Taken together, these results indicate no change in peripheral nerve sprouting, apparent inflammation nor bone alterations following 10nmol LPC16:0 joint injections.

### The *in vivo* pronociceptive effects of LPC16:0 are largely dependent on ASIC3 in a sex-independent manner

We had shown previously that LPC16:0 activate ASIC3 *in vitro*^6^, and thus investigated here whether this channel participates to the establishment and/or maintenance of pain and anxiety-like behaviors induced by intra-articular LPC16:0 injections using wild-type littermates (ASIC3^+/+^) and ASIC3 knockout (ASIC3^−/−^) mice (**fig. 4 and supplementary fig. 4**). Intra-articular administrations of vehicle did not change mechanical paw withdrawal threshold in male or female ASIC3^+/+^ mice, which remained similar to baseline throughout the experiment (**fig. 4a–b**). In contrast, LPC16:0 injections induced long-lasting mechanical allodynia up to day 28 and day 26 in male and female ASIC3^+/+^ mice, respectively, indicating the absence of sex dimorphism (**fig. 4a–b**). The peak intensity of LPC-induced mechanical allodynia was significantly reduced by around 50-60% in both male and female ASIC3^−/−^ mice, and the duration of the effects was shorter with paw withdrawal thresholds back to baseline levels before the end of the experiment (**fig. 4a–b**). Both male and female ASIC3^+/+^ mice also developed significant and long-lasting thermal hyperalgesia following LPC16:0 injections, which was drastically reduced in ASIC3^−/−^ mice of both sexes (**supplementary fig. 4a-b**). In the open field test, LPC16:0 did not alter the locomotor activity of male or female ASIC3^+/+^ and ASIC3^−/−^ mice compared to control groups, as illustrated by a similar distance travelled between groups (**supplementary fig. 4c,d**). In the elevated plus maze test, the time spent in the open arms was significantly reduced by 53% and 60% in LPC16:0-injected male and female ASIC3^+/+^ mice, respectively, compared to vehicle-injected ASIC3^+/+^ mice (**fig. 4c,d**), indicating no sex dimorphism in the development of anxiety-like behaviors associated with LPC16:0. Such reduction in the time spent in the open arm was not observed in male and female ASIC3^−/−^ mice (**fig. 4c,d**). In the marble burying test, the number of buried marbles was significantly higher in both male and female ASIC3^+/+^ mice injected with LPC16:0, compared to ASIC3^+/+^ control mice (**fig. 4e,f**). This increased number of buried marble was not observed in LPC16:0-injected male or female ASIC3^−/−^ mice (**fig. 4e,f**). Thus, the development of LPC16:0-induced pain-like behaviors were significantly reduced and the associated anxiety-like behaviors were absent in ASIC3^−/−^ mice compared to ASIC3^+/+^ mice of both sexes.

**Figure 4:**
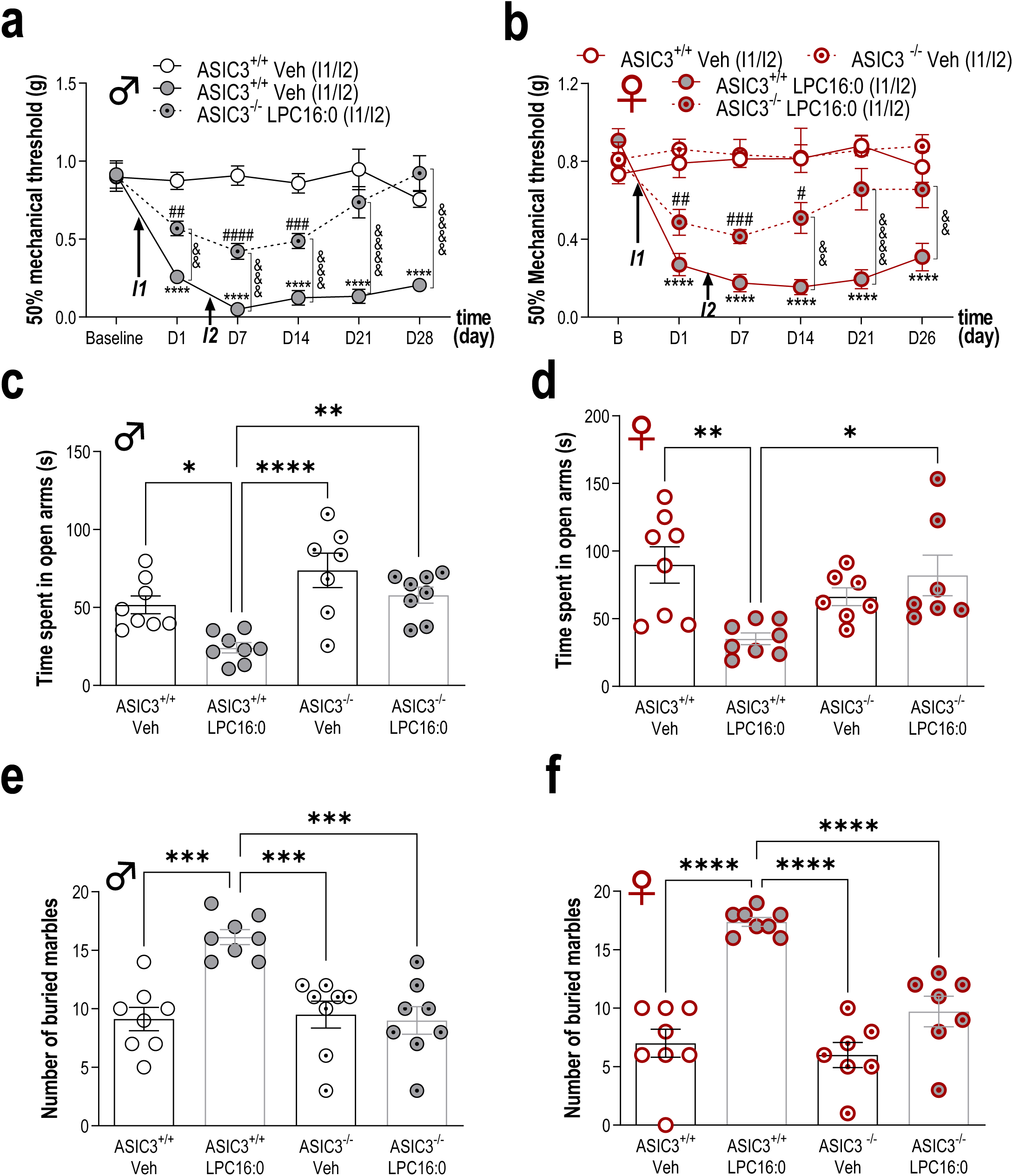
ASIC3 is crucial for the development of both pain and anxiety-like behaviors induced by ankle LPC16:0 injections in male and female mice. Time course effect of intra-articular ankle injections of LPC16:0 (10 nmol) or vehicle in male (**a**) or female (**b**) ASIC3^+/+^ and ASIC3^−/−^ mice. Mechanical paw withdrawal threshold was assessed using the up and down method with von Frey filaments from D1 to D28, and D1 to D26, for male and female mice, respectively (n=7-8 mice per group, \*\*\*\**p*<0.0001 for ASIC3^+/+^ Veh *vs.* ASIC3^+/+^ LPC16:0; #p<0.05, ^##^*p*<0.01, ^###^*p*<0.001 and ^####^*p*<0.0001 for ASIC3^−/−^ LPC16:0 *vs.* ASIC3^+/+^ Veh 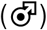 or ASIC3^−/−^ Veh 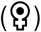; ^&&^*p*<0.01 and ^&&&&^*p*<0.0001 for ASIC3^+/+^ LPC16:0 *vs.* ASIC3^−/−^ LPC16:0; Two-way ANOVA followed by Tukey’s *post hoc* tests). **c-f**, Effect of intra-articular ankle injections of LPC16:0 (10 nmol) or vehicle on the time spent in the open arm in the elevated plus maze test (**c-d)**, on the number of buried marble in the marble burying test (**e-f)**, in male (**c-e**) or female (**d-f**) ASIC3^+/+^ and ASIC3^−/−^ mice. The elevated plus maze test lasted 5 min and the time spent in the open arm was automatically calculated using Ethovision XT 13 (Noldus). The marble burying test lasted 30 min and was analyzed by a blind experimenter (n=7-8 mice per group, \**p*<0.05, \*\**p*<0.01 and \*\*\*\**p*<0.0001, one-way ANOVA followed by Tukey’s *post hoc* tests).

### LPC16:0 induces an ASIC3-dependent membrane depolarization in DRG neurons

Extracellular application of LPC16:0 in HEK293 cells transfected with rat or human ASIC3 channels induced a progressive and non-inactivating inward current at resting pH7.4, *i.e.* without any extracellular pH changes (**supplementary fig. 5a, c and d)**. Conversely, no significant LPC-activated currents were observed in ASIC1a-transfected cells, nor in non-transfected ones (**supplementary fig. 5b, d**). Furthermore, a LPC16:0-induced current was observed in primary cultured mouse DRG neurons (**fig. 5**). This current had a significantly higher density in wild-type neurons that expressed a pH6.6-induced native ASIC current (WT ASIC+) compared to neurons without native ASIC current (WT ASIC-), and to neurons isolated from ASIC3 knockout mice, expressing (KO ASIC+) or not-expressing (KO ASIC-) native ASIC currents (**fig. 5a–c**). Therefore, ASIC3 contributed largely to the LPC16:0-induced currents in mouse DRG neurons. A small ASIC3-independent current was still observed, suggesting an effect of LPC16:0 on other native ion channels, as expected^5,15,16^. Finally, the neuronal depolarization associated to a 30-sec application of LPC16:0 was significantly higher in WT ASIC+ neurons than in WT ASIC-, KO ASIC+ or KO ASIC- neurons, supporting a leading role of ASIC3 in the excitatory effect of LPC16:0 on DRG neuron membrane potential (**fig. 5d**).

**Figure 5:**
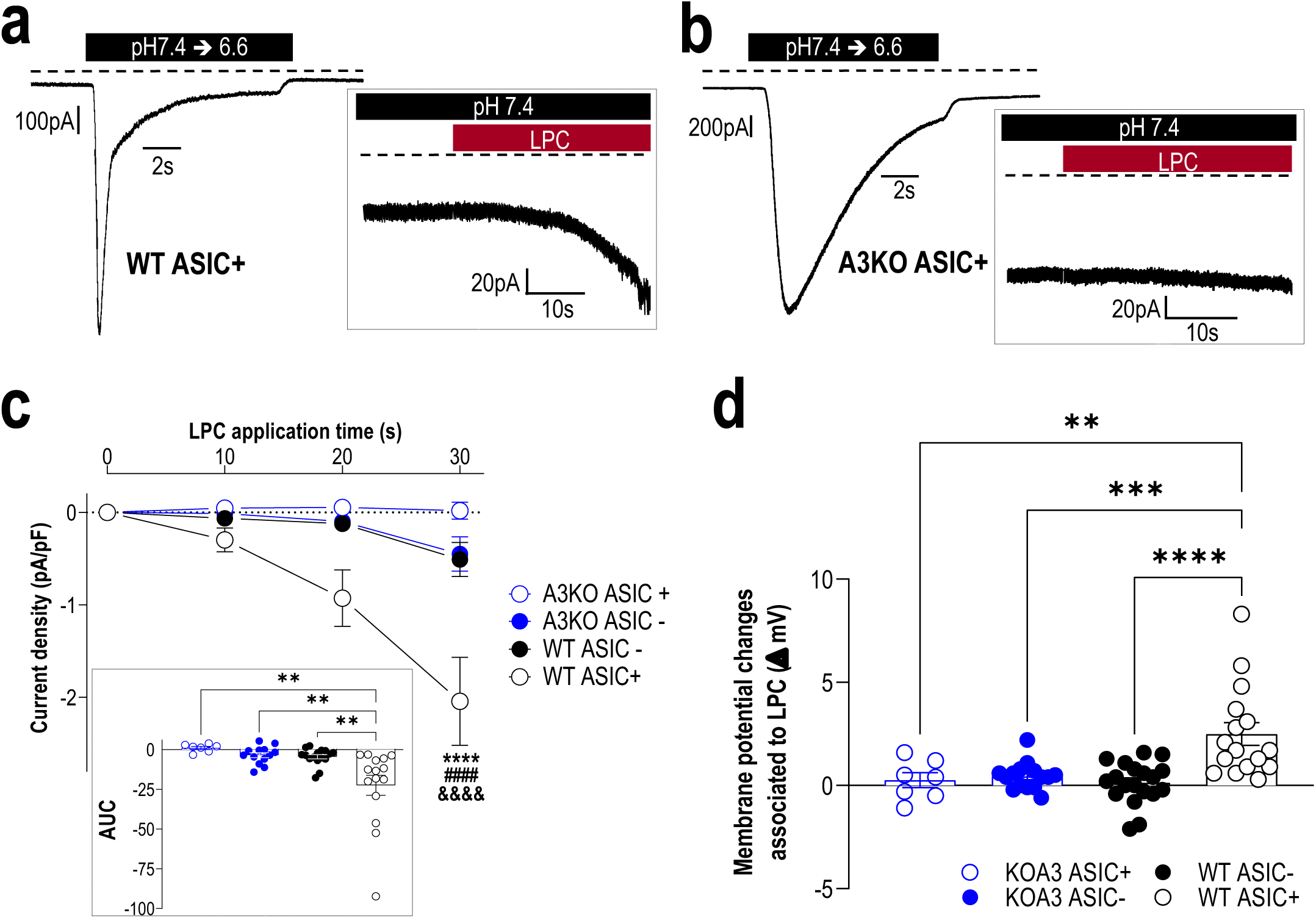
LPC16:0 induces a non-inactivating, ASIC3-dependent current associated with membrane depolarization in mouse DRG neurons. **a-b**, Native pH6.6-evoked ASIC currents recorded at −80 mV in DRG neurons from wild-type (WT ASIC+, **a**) and ASIC3 knockout (KO ASIC+, **b**) mice. *Insets* show the effect of LPC16:0 (5μM) applied for 30 seconds on the same neurons. **c**, Analysis of the current densities measured after 10, 20 and 30 second-applications of LPC16:0 onto WT and ASIC3 KO DRG neurons expressing (ASIC+) or not expressing (ASIC-) a pH6.6-evoked ASIC current (see methods, n=7-16; \*\*\*\**p*<0.0001 compared to WT ASIC-, ^####^*p*<0.0001 compared to KO ASIC+ and ^&&&&^*p*<0.0001 compared to KO ASIC-; Three-way ANOVA followed by a Tukey’s multiple comparison test). *Inset* shows the net AUC calculated over 30 second period from the baseline fixed at 0 (see methods, n=7-16, \*\**p*<0.01, one-way ANOVA followed by a Tukey’s multiple comparison test). **d**, Membrane potential changes associated with 30 second-application of LPC16:0 on WT and ASIC3 KO DRG neurons (n=7-19; \*\**p*<0.01, \*\*\**p*<0.001 and \*\*\*\**p*<0.0001, one-way ANOVA followed by a Tukey’s multiple comparison test).

### LPC16:0 injections into mouse knee joints also produce persistent pain-like behaviors, which are associated to spinal neuron sensitization

LPC16:0 was injected into mouse knee joint to determine if the results obtained with the ankle can be extended to other joints (**fig. 6 and supplementary fig. 6**). Pain and anxiety-like behaviors were assessed at different time points after intra-articular knee injections of LPC16:0 at two different doses, *i.e.*, 10 nmol as for the ankle, and 20 nmol (**fig. 6a**). Mice injected in the knee with LPC16:0 developed a dose-dependent mechanical allodynia assessed with the von Frey test (**fig. 6b**), significant weight bearing deficit and thermal hyperalgesia (**supplementary fig. 6a-b)**, associated with anxiety-like behaviors (**supplementary fig. 6c-d**) not due to locomotor impairment (**supplementary fig. 6e**). At these doses, LPC16:0 effect was not due to potential nerve demyelination^27,28^, since we did not observe any demyelination of the saphenous nerve following 20nmol LPC16:0 knee injections, as demonstrated by the similar G-ratio between vehicle and LPC16:0 groups (**supplementary fig. 3f**). When mechanical withdrawal thresholds were assessed with the dynamic plantar aesthesiometer test, LPC16:0 injections (20 nmol) into mouse knees also induced long-lasting allodynia (**fig. 6c**) that was clearly secondary to the injection site. This effect was only ipsilateral (**supplementary fig. 6g**) and observed in both male and female wild-type (WT) mice with comparable kinetics, amplitudes and durations (**fig. 6c**). As for the ankle joint, LPC16:0 knee injections-induced pain was significantly reduced in both male and female ASIC3^−/−^ mice compared to WT mice, with a more pronounced reduction here (**fig. 6d**). This long-lasting pain state was also significantly reduced in male WT mice when APETx2, an ASIC3 pharmacological blocker^6,29^, was co-injected with LPC16:0 either at the first or the second administration, further supporting an effect primarily mediated through peripheral ASIC3 activation (**fig. 6e**). Finally, two LPC16:0 intra-articular injections were necessary, as for the ankle model (**fig. 2b**), to induce a persistent pain state, *i,e*., a single knee injection only produced transient mechanical allodynia (**supplementary fig. 6h**). These data supported a similar LPC-induced ASIC3-dependent persistent pain state upon two intra-articular injections of LPC16:0 in either knee or ankle joints, which suggested a central sensitization process. The activity of spinal dorsal horn neurons was therefore recorded *in vivo* in painful WT mice following LPC16:0 (20 nmol) knee injections (day≥30). Recordings were obtained from spinal dorsal horn neurons receiving either non-noxious sensory inputs (low threshold neurons, LTNs) or noxious nociceptive inputs (high threshold neurons, HTNs), with receptive fields located in mice hindpaws. Brushing-evoked activity of ipsilateral LTNs, *i.e.,* the mean number of emitted action potentials (APs), was not modified in LPC16:0-treated mice compared to vehicle or to contralateral LTNs (**fig. 6f**). Conversely, the evoked activities of HTNs in response to both pinch (**fig. 6g**) or von Frey filaments (**fig. 6h**) were significantly enhanced in mice injected with LPC16:0. Ipsilateral HTNs of LPC16:0-treated mice emitted on average twice as much APs in response to pinch as ipsilateral HTNs from vehicle-treated mice or contralateral HTNs (**fig. 6g**). Moreover, ipsilateral HTNs of LPC16:0-treated mice clearly displayed mechanical sensitization in response to the application of increasing von Frey filaments, with a response curve significantly shifted to the left compared to contralateral HTNs or to ipsilateral HTNs from vehicle-treated mice (**fig. 6h**). *In vivo* recordings of spinal dorsal horn neurons clearly demonstrated a sensitization of spinal high threshold, but not low threshold neurons induced by LPC16:0 knee injections, which was correlated with the persistent secondary allodynia observed.

**Figure 6:**
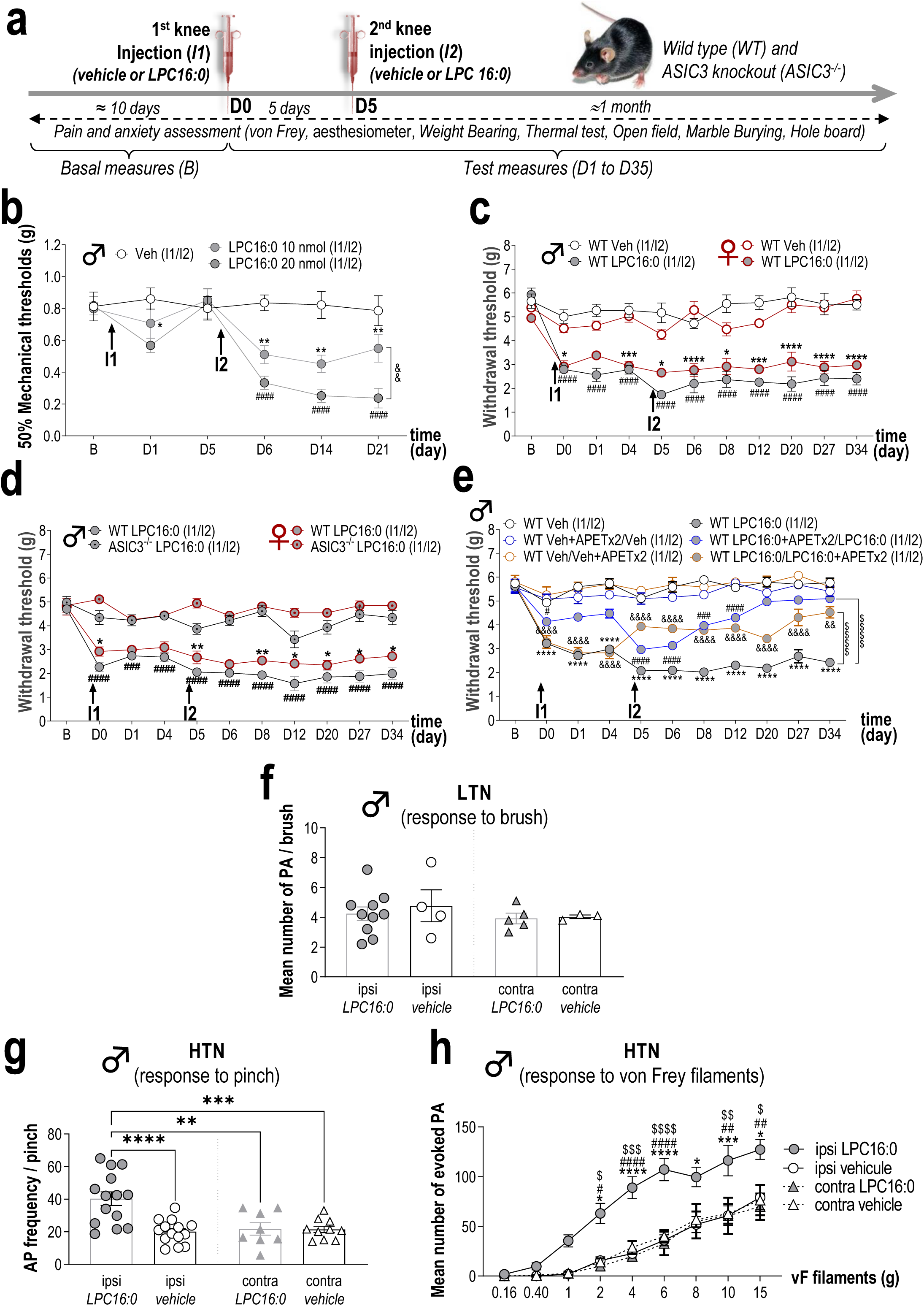
LPC knee injections generate persistent secondary mechanical allodynia associated with sensitization of spinal dorsal horn neuron activity. **a**, Timeline of the experimental procedures used. **b**, Effect of intra-articular knee administrations of LPC16:0 (10 and 20 nmol), or vehicle (Veh) on mechanical allodynia in the ipsilateral paw of male mice. Mechanical paw withdrawal threshold was assessed using the up and down method with von Frey filament from D1 to D21. Results are expressed as 50% mechanical thresholds (n=8 mice per group; \**p*<0.05 and \*\**p*<0.01 for LPC 16:0 10 nmol *vs.* Veh, ^####^*p*<0.0001 for LPC16:0 20 nmol *vs.* Veh and ^&&^p<0.01 for LPC16:0 20 nmol *vs.* LPC16:0 10 nmol; Two-way ANOVA followed by a Tukey’s *post hoc* test). **c**, Ipsilateral paw withdrawal thresholds (PWTs) of female and male WT mice injected twice in the knees with LPC16:0 (20 nmol) or vehicle (Veh). PWTs were assessed using a dynamic plantar aesthesiometer (n=6 mice per group; \**p*<0.05, \*\*\**p*<0.001 and \*\*\*\**p*<0.0001 for WT female Veh *vs.* WT female LPC16:0; ^####^*p*<0.0001 for WT male Veh *vs.* WT male LPC16:0; Three-way ANOVA test followed by a Tukey’s multiple comparison test). **d**, Ipsilateral PWTs of female and male, WT and ASIC3^−/−^ mice injected twice with LPC16:0 (n=5-18 mice per group; \**p*<0.05, and \*\**p*<0.01 for female WT LPC16:0 *vs.* ASIC3^−/−^ LPC16:0; ^###^*p*<0.001 and ^####^*p*<0.0001 for male WT LPC16:0 *vs.* ASIC3^−/−^ LPC16:0; Three-way ANOVA test followed by a Tukey’s multiple comparison test). **e**, Ipsilateral PWTs of male WT mice injected twice with LPC16:0 or vehicle, with or without co-injection of APETx2 (0.1 nmol) at the first or the second injection (n=6-12 per group; \*\*\*\**p*<0.0001 for WT Veh *vs.* WT LPC16:0; ^#^*p*<0.05, ^###^*p*<0.01 and ^####^*p*<0.0001 for WT Veh+APETx2/Veh *vs.* WT LPC16:0+APETx2/LPC16:0; ^&&^*p*<0.01 and ^&&&&^*p*<0.0001 for WT Veh/Veh+APETx2 *vs.* WT LPC16:0/LPC16:0+APETx2; Two-way ANOVA test followed by a Tukey’s multiple comparison test; ^$$$$^*p*<0.0001 for main effects on curves, two-way ANOVA test followed by a Tukey’s multiple comparison test). **f**, Mean number of action potentials (APs) emitted by spinal low threshold neurons (LTNs) following light brushing of their receptive fields, located on the plantar surface. Spinal LTN evoked-activities were not different whatever they were recorded ipsilaterally or contralaterally to the knees injected twice with LPC16:0 or vehicle (n=3-10 LTNs, no significant differences, Kruskal-Wallis test followed by a Dunn’s *post hoc* test). **g**, Frequencies of APs emitted by spinal HTNs following pinching (n=8 to 14 HTNs; \*\**p*<0.01, \*\*\**p*<0.001 and \*\*\*\**p*<0.0001, one-way ANOVA followed by a Tukey’s multiple comparison test). **h**, Mean number of APs evoked by spinal HTNs following application of von Frey filaments of increasing strength (n= 8 to 14 HTNs; \**p*<0.05, \*\*\**p*<0.001 and \*\*\*\**p*<0.0001 for ipsi LPC16:0 *vs*. ipsi vehicle; ^#^*p*<0.05, ^##^*p*<0.01 and ^####^*p*<0.0001 for ipsi LPC16:0 *vs*. contra LPC16:0; ^$^*p*<0.05, ^$$^*p*<0.01, ^$$$^*p*<0.001 and ^$$$$^*p*<0.0001 for ipsi LPC16:0 *vs*. contra vehicle; Three-way ANOVA test followed by a Tukey’s multiple comparison test).

## Discussion

We identify LPC16:0 as the major lysophosphatidyl-choline species in human synovial fluids and a critical trigger of chronic joint pain through peripheral ASIC3 channels activation in a mouse preclinical model. An elevated content of LPC16:0 was observed in the knee synovial fluids of a first cohort of patients with OA, as compared to controls, and this content was correlated to knee and global pain (VAS knee and VAS global), regardless of age, gender, BMI and IL-6 synovial fluid levels. Correlation with knee osteoarthritis outcome score (KOOS) almost reach statistical significance (*p*=0.060). This absence of a statistically significant correlation can be explained by the more holistic nature of the patient’s overall OA experience assessed by this questionnaire, where excess nociception, emotions, the impact of joint damage on the patient’s quality of life and personality are intertwined. A specific increase of LPC16:0 content was further confirmed in the synovial fluids of a second cohort of patient with different joint diseases, including OA but also other pathologies such as rheumatoid arthritis (RA). These results suggest that LPC is probably not the hallmark of a particular joint pathology. Previous works also reported an increase of plasma and/or synovial LPC contents in OA and rheumatoid arthritis (RA) patients^17,30,31^, but without informations on its possible contribution to chronic joint pain.

We demonstrate that ankle or knee intra-articular administrations of LPC16:0 in mice lead to the development of chronic pain-related behaviors as well as anxiety-like behaviors, independently of LPC-to-LPA conversion by autotaxin. Using this new preclinical model of persistent joint pain, our data support a direct role of LPC16:0 as a triggering factor of pain-like behaviors in mice, with the potential to also have pain-related roles in humans. The persistent pain state in mice did not seem to be associated with nerve sprouting, demyelination or inflammatory processes, since no changes in CGRP joint immunoreactivity, nerve fiber G ratio, *in vivo* metalloprotease activity, nor expression of TNF and IL-6 were observed in joints following LPC16:0 administrations. Conversely, these cytokines appeared upregulated in CFA-injected joints, as expected. Hung and colleagues^18^ also failed to observe any inflammatory signs following intramuscular administration of LPC16:0 in mouse muscle. Because bone alteration/remodeling has been suggested to contribute to chronic joint pain in rheumatic diseases such as OA^32–34^ and RA^35^, the Cathepsin K activity as well as joint expression of TRAP and RANK-L has been investigated in our model. No changes of these different markers were observed, indicating that LPC16.0 did not lead to bone alterations, and suggesting that it may rather be its consequence.

On the other hand, our work reveals that the pronociceptive effects of LPC16:0 are dependent on ASIC3 channels expressed in nociceptors, which play a sex-independent role in this process. Both pain and anxiety-like behaviors were indeed significantly reduced in both male and female ASIC3 KO mice, or following local pharmacological inhibition of these channels. The contribution of ASIC3 in the development of pain-like behaviors evoked by LPC16:0 appeared to be more significant following injections into the knee than into the ankle of mice. Indeed, ASIC3 knockout mice were almost fully protected against LPC16:0-induced pain following knee injections, whereas pain-related behaviors still remained, although largely decreased, in case of ankle injections. Such a difference might illustrated a predominant role of ASIC3 in the development of secondary pain, as proposed elsewhere^22,25^. Since the pain measurements were taken at the plantar level, they more likely reflected a mixture of primary and secondary pain in the case of ankle injections, whereas they were clearly associated to secondary pain following knee injections. ASIC3, which was constitutively activated by LPC16:0 in cultured DRG neurons, induced a sustained membrane depolarization similar to the ASIC3-dependent depolarization already shown to sensitize these neurons^36^. It is thus very likely that the two consecutive intra-articular injections of LPC16:0 drive a nociceptive ASIC3-dependent input leading to a persistent pain state originating from joints. ASIC3 has been already associated to long-lasting pain states following acid injections in rodent joints^24^ and muscles^25^ and it has been more recently proposed to participate in pain behaviors induced by LPC16:0 muscle injections in mice^18^. The stimulation of ASIC3 channels in nerve endings of deep tissues, such as muscles or joints, is thus a key step for the development of persistent pain state.

This persistent pain state generated by LPC16:0 requires two injections similarly to what was described for intraplantar cytokine (IL-6, TNF) injections combined with subsequent PGE2 administration, and for repeated acid injections into muscle or joint^22,24,25,37^. Moreover, it is not modality specific since LPC16:0-treated mice displayed primary and secondary mechanical allodynia, thermal hyperalgesia and weight bearing deficit, together with anxiety-like behaviors. The development of secondary allodynia after LPC16:0 joint administrations suggests potential central sensitization processes. Indeed, LPC16:0 sensitized dorsal horn neurons, most probably through its effect on ASIC3 expressed in joint nociceptor endings, as demonstrated by the enhanced output of spinal high threshold nociceptive, but not low threshold, sensory neurons. Repeated intramuscular injections of LPC16:0 induced an increased expression of c-fos and pERK in dorsal horn neurons, also consistent with central sensitization^18^. Several clinical studies provide evidences for the presence of central sensitization in chronic OA pain patients^38^. The elevated level of LPC, especially LPC16:0, observed in the synovial fluid of OA patients could explain why they still suffered from chronic joint pain despite well controlled inflammation and possibly even in the absence of it, as suggested by the lack of correlation observed between LPC16:0 and IL-6 levels in their synovial fluids. Since LPC16:0 administrations did not apparently alter the injected joints, this might also support the lack of clear correlation between joint damage and pain symptoms in OA^39,40^.

To conclude, LPC16:0 could be involved in joint pain from different etiologies of inflammatory and non-inflammatory origins, and more generally across rheumatic musculoskeletal diseases. However, it remains to be demonstrated whether an elevated level of LPC16:0 can be used as an objective biomarker of chronic joint pain across different etiologies. Our data bring both human and rodent evidences for a crucial role of this lipid in chronic joint pain, and identify LPC-activated ASIC3 channels as very attractive targets for chronic joint pain management.

## Methods

### Patient cohorts

#### Cohort of patients with osteoarthritis (OA, first cohort)

Thirty-five patients with knee OA (16 women and 19 men, average age 64.8 years, range 49‐73 years) participated. This study was approved by the Regional Ethical Review Board in Stockholm, Sweden (reference number 2011/2036-31-1, 2012/2006-32) and followed the guidelines of the Declaration of Helsinki. All patients were recruited consecutively from the waiting list for total knee replacement at Ortho Center, Upplands Väsby, Sweden. The patient inclusion criteria were 25‐75 years of age, radiologically verified knee OA and the presence of knee pain as the dominant pain symptom and a motivation for surgery. The patients were excluded if they suffered from chronic pain due to causes other than knee OA (e.g., fibromyalgia, degenerative disc disease, disc herniation, inflammatory rheumatic disease or neurologic disease) or in case of previous knee surgery planned for total knee replacement. Information regarding medication was collected from all patients. Eight patients were taking analgesics (3 codeine, 2 tramadol, 2 buprenorphin plaster, 1 Ketobemidone), 12 were taking acetaminophen and 16 had previously been taking nonsteroidal anti-inflammatory drugs (NSAIDs) at demand, however these had been stopped 14 days before the surgical procedure. All patients received 2 g acetaminophen (paracetamol) and 10 mg oxycodone orally as premedication before surgery. All participants were informed about the study procedure, and written consent was obtained. Knee joint synovial fluid was collected during surgery and immediately frozen at −80°C for future analysis.

#### Cohort of patients with different joint diseases (second cohort)

Fifty patients suffering from different painful joint diseases (32 women and 18 men, average age 68.6 years, range 26-94 years), recruited in the Rheumatology Department of Nice University Hospital, France, participated and were distributed as follow: rheumatoid arthritis (RA, n=6), chondrocalcinosis (CCA, n=12), spondyloarthritis (SPA, n=5), psoriatic arthritis (PA, n=4), gout (n=5) and OA (n=18). All subjects provided informed consent before inclusion and the study was approved by the Nice University Institutional Review Board for Research on Human Subjects. The study has been conducted in accordance with the French national regulations regarding patient consent and ethical review. The study was registered in the ClinicalTrials.gov protocol registration system (NCT 01867840). All samples were obtained from patients with acute knee joint effusion requiring joint puncture for diagnosis and/or treatment. The synovial fluids remaining after biological analysis for patients’ care were included in the present study and frozen at −80°C for future analysis.

#### Post mortem controls

Ten post-mortem (PM) subjects with no history of knee or hip OA or inflammatory rheumatic diseases were included as controls. Synovial fluid from the knee joint was collected during an autopsy procedure and immediately frozen at −80°C for future analysis.

#### Questionnaires

Only OA patients from the first cohort completed questionnaires and scales within a week before surgery. Global pain intensity at the day of examination (VAS global) and pain in the affected knee (VAS knee) were all scored using 100 mm VAS with 0 indicating “no pain” and 100 indicating “the worst imaginable pain”. The severity of patient-reported symptoms was assessed by Knee injury and Osteoarthritis Outcome Score (KOOS), which consists of 5 subscales: a) pain, b) other symptoms, c) activity in daily living, d) function in sport and recreation, and e) knee related quality of life^41,42^. Each KOOS subscale contains questions scored from 0 to 4 and the average score of all 5 KOOS subscales was calculated and used for the analysis.

#### Patient demographic data and LPC concentrations

The demographic data (age, sex, BMI) and LPC concentrations found in patients synovial fluids form the first (OA patients) and the second cohort, as well as in PM controls, are reported in **supplementary tables 1 and 3**. Part of the data of the first cohort (age, sex, BMI, VAS global, VAS knee, KOOS assessment and IL-6 levels in the knee synovial fluids) have also been reported elsewhere^43^.

### Lipodomic analysis of patient samples

#### Chemical and Lipid Standards

Chloroform (CHCl_3_), Methanol (CH_3_OH) and Formic acid (HCOOH) were obtained from Sigma Aldrich (Saint Quentitn Fallavier, France). Water (H_2_O) used for lipid extraction was from Milli-Q quality. LPC standards were purchased from Avanti Polar Lipids *via* Sigma Aldrich, and then prepared at the appropriate concentration and stored at −20°C.

#### Lipid extraction

All patients’ samples were processed in the same manner by the same laboratory as follow. Extraction of lipids from human synovial fluid (HSF) samples was adapted from the Folch method^44^. Fifty microliters of each patient sample were diluted with water to a final volume of 1 ml. The diluted samples were then transferred into glass tubes (Pyrex Labware) and vortexed for 1 min. Lipids were extracted using 4 ml Chloroform (CHCl_3_)/methanol (CH_3_OH) (2:1, v/v) and shaken with an orbital shaker (IKA® VX® basic Vibrax®, Sigma-Aldrich) at 1,500 rpm for 2 h at room temperature. After centrifugation for 10 min with a swing-out centrifuge at 410 g, aqueous phases were eliminated, and lipid-containing organic phases were supplemented with 1 ml of a 4:1 (v/v) 2N KCl/CH_3_OH solution. Samples were shaken for 10 min at 1,500 rpm, centrifuged for 5 min and the upper aqueous phases were eliminated. The resulting organic phases were complemented with 1 ml of a 3:48:47 (v/v/v) CHCl_3_/CH_3_OH/H_2_O solution, shaken for 10 min and centrifuged for 5 min. The aqueous phases were eliminated and the organic phases containing whole lipids were transferred into new glass tubes. Lipid extracts were then evaporated until dryness at 60°C under a stream of nitrogen, re-dissolved in 500 μl CHCl_3_/CH_3_OH (1:2, v/v) and stored at −20°C until further analysis.

#### Mass spectrometry analysis of lysophosphatidyl-choline (LPC)

Lipid extracts were diluted once in CHCl_3_/CH_3_OH (1:2, v/v) before addition of 1 % (v/v) formic acid. An optimized quantity of the internal standard LPC13:0 (0.1 μg of LPC13:0 for 100 μl of lipid extract) was added to each sample in order to quantify LPC species. Diluted lipid extracts were analyzed in the positive ion mode by direct infusion on a SYNAPT^™^ G2 High Definition Mass Spectrometer (HDMS; Waters Corporation, Milford, Massachusetts, USA) equipped with an Electrospray Ionization Source (ESI). The flow rate was 5 μl/min. All High Resolution full scan Mass Spectrometry (HR MS) experiments were acquired in profile mode over 1min with a normal dynamic range from 300 to 1,200 m/z. Ionization conditions in positive ion mode have been optimized and all LPC species, including the LPC internal standard, were detected as protonated ions [M+H]^+^.

Identification of LPC species was based on their exact masses in the full scan spectrum and on tandem mass spectrometry experiments (MS/MS) in the positive ion mode. MS/MS fragmentation was performed by collision-induced-dissociation (CID), which notably allowed to obtain structural information about LPC species by identifying the choline polar head group (characteristic and prominent fragment ion for all LPC species with *m/z* 184).

LPC species were quantified by normalizing the intensity of the protonated ion [M+H]^+^ corresponding to each LPC individual species in the full scan spectrum to the intensity of the internal standard LPC 13:0 [M+H]^+^, and multiplying by the amount of the internal LPC standard. In addition, corrections were applied to the data for isotopic overlap. Total LPC amount was calculated by summing the quantities of all individual LPC species. All spectra were recorded with MassLynx software© (Version 4.1, Waters). Data processing for high resolution full scan experiments was carried out with the help of ALEX Software^45^.

### Animals and behavioral experiments

#### Animals

Experiments were performed on adult male and/or female C57Bl6J wild‐type mice (WT, Janvier labs) as well as ASIC3 knockout (ASIC3^−/−^) and wild‐type littermates (ASIC3^+/+^), breed in animal facilities of IPMC and UCA Medicine School with agreements n°C061525 and n°C63115.15, respectively. Animals were kept with a 12‐h light/dark cycle with *ad libitum* access to food and water and were acclimated to housing and husbandry conditions for at least 1 week before experiments. All experiments followed guidelines of the International Association for the Study of Pain^46^. All the protocols used were approved by the local ethical committees (CIEPAL-Azur and CEMEA-Auvergne) and the French government (agreements n°02595.02, APAFIS#13499, APAFIS#17387) in accordance with European Communities Council Directive for the care of laboratory animals (86/609/EEC).

#### Joint pain models

Monoarthritis (MoAr) was induced by a single intra-articular injection of 12 μl of Complete Freund Adjuvant (CFA, heat-killed *Mycobacterium butyricum* 2 mg/ml, Sigma) under isoflurane (2.5%) anesthesia and was used as a positive control. Concerning the LPC-induced joint pain model, mice received two consecutive intra-articular injections of saline solutions containing either LPC16:0 (10 or 20 nmol, Avanti polar lipds, Coger, France) or vehicle (EtOH 2-4%), five days apart, within the ankle or knee joints (10μl) under isoflurane (2.5%) anesthesia. The mechanical (von Frey test) and heat (thermal test) pain thresholds, as well as the weight bearing between hindpaws (static weight bearing test) were measured before and after LPC/CFA/vehicle injections over a month period (see **figs. 2a and 6a** for schematic protocol of experiments).

#### Pharmacological experiments

The autotaxin inhibitor S32826 (Biotechne, France), was injected in mice ankles (10 nmol) either at the time of the first LPC16:0 injection, or 5 days after the second LPC16:0 injection. When S32826 was co-injected with the first LPC16:0 administration, mechanical allodynia was assessed from 30 min to 240 min post-injection. For experiments where S32826 was administrated 5 days after the second LPC16:0 injection, the mechanical allodynia was assessed 30 min and 60 min post-injection. APETx2 (Smartox, France), the ASIC3 pharmacological inhibitor^29^, was co-injected in mice knees together with LPC16:0 (0.1 nmol APET×2 + 20 nmol LPC16:0), either at the first or the second intra-articular administration of LPC16:0.

#### Mechanical sensitivity

Mechanical sensitivity was evaluated by performing von Frey tests. Two types of von Frey tests were used, the classical up-down method of Dixon^47^ modified by Chaplan^48^ with von Frey filaments (Bioseb, France) and the dynamic plantar aesthesiometer (Ugo Basile, Italy). For both methods, freely moving animals were placed in individual plastic boxes on a wire mesh surface, so that von Frey mechanical stimuli could be applied to the plantar surfaces of their hindpaws. For up-down von Frey experiments, animals followed an habituation period before a set of calibrated von Frey filaments ranging from 0.02 to 1.4g was applied to the plantar surface of each hindpaw, alternatively or only on the ipsilateral hindpaw, until withdrawal or licking of the paw to the stimulus. The 50% paw mechanical threshold was evaluated before (Baseline) and at different time points after the first and second LPC16:0 or vehicle injections, or after the CFA injection. For experiments using the dynamic plantar aesthesiometer, a ramp of force was applied through a single filament (up to 7.5g in 10s), and paw withdrawal thresholds (g) were measured in duplicate. A mean withdrawal threshold was then calculated for each animal hindpaw. Three baseline measures were made prior LPC16:0 or vehicle intra-articular injections, the third baseline measure served as the reference mechanical withdrawal threshold for statistical analyses.

#### Heat sensitivity

Heat sensitivity in mice was assessed by the paw immersion test using a thermo-regulated water bath maintained at 46.0 ± 0.2 °C. Mice were maintained under a soft cloth except the ipsilateral hindpaw, which was immersed in water until withdrawal (a cut-off of 30s was used to avoid any potential tissue damage).

#### Weight bearing experiments

Static weight bearing (SWB) apparatus (Bioseb, France) was used to evaluate spontaneous weight distribution between hindpaws, which could be considered as a measure of ongoing pain. Following two training sessions to habituate animals, mice were placed in a plastic glass enclosure so that each hindpaw rested on separate transducer pads. Once mice were settled in correct position, the force exerted by each hindpaw was measured over a period of 5s. Weight-bearing differences were recorded as an average of four/five trials and expressed as contralateral/ipsilateral ratio. The weight distribution was assessed before (Baseline) and at different time points following vehicle, CFA or LPC injections.

#### Assessment of anxiety-like behaviors

##### Open field test

Mice were individually placed in the middle of an open field (50 × 50 × 50 cm) under dim light conditions (30 lux). The distance travelled and the velocity were automatically calculated during the 5 min testing period as well as the time spent in the center for each mouse (Ethovision XT 13, Noldus).

##### Elevated Plus Maze

The elevated plus maze (EPM) consists of four arms, two opposite open arms (37.5 × 5 cm) and two opposite closed arms (37,5 × 5 cm with 20 cm high walls), joined by a common central platform (5 × 5 cm), subjected to an equal illumination (30 lux). The maze was elevated to 60 cm above the floor. Each mouse was placed randomly into the center of the EPM facing an open arm. The time spent into the open arms, considered when head, gravity center and tail points were located within the arm, was automatically recorded for 5 min and calculated for each animal (Ethovision XT 13, Noldus).

##### Marble burying test

Briefly, mice were individually placed in the experimental cage (42.5 × 27.6 × 15.3 cm) containing 20 marbles (4 lines of 5 equidistant marbles) disposed on the top of 5 cm thickness litter. Animals were left undisturbed during 30 min in a quiet room. At the end, mice were removed and the numbers of buried marbles were quantified. A marble was considered as buried when at least 50% of its surface was covered with litter.

##### Hole board test

The hole board test consists of a board (39.5 × 39.5 cm) with 16 equidistant holes, 3 cm in diameter, equally distributed throughout the platform and placed 70 cm above the floor. Mice were individually placed randomly on one corner of the board facing away the experimenter and videotaped. The number of head dips in the holes was quantified for 5 min by a blind experimenter.

### Immunohistochemistry on tibiotarsal joints and amygdala

At day 28, four and five mice of the vehicle and LPC16:0 10 nmol groups, respectively, were terminally anaesthetized using a mixture of Ketamine/Xylazine and quickly perfused transcardially with saline followed by 4% paraformaldehyde (PFA). Ipsilateral tibiotarsal joints were excised, post fixed in 4% PFA in phosphate buffer (0.1 M, pH 7.4) for 48h at 4°C, and decalcified in 10% ethylenediaminetetraacetic acid (EDTA, pH 7.6, Sigma) during two weeks at 4°C as previously described^49^. Brain was also excised and post fixed in 4% PFA in phosphate buffer (0.1 M, pH 7.4) for 24h at 4°C. After cryoprotection (PB-Sucrose 30%) for at least 48h, samples were included in tissue freezing medium (O.C.T.).

Twenty μm cryostat thick frozen sections of joints were processed, mounted on Superfrost^®^ slides, blocked with PBS, BSA 1% and, incubated with primary antibodies in PBS + Triton 0.2% overnight at room temperature following three washes in PBS. Monocytes, macrophages and osteoclasts lineage was labelled with a rat antibody against myeloid protein CD68 (1:1,000, AbD Serotec, #MCA1957) and peptide-rich sensory nerve fibres with a rabbit anti-CGRP (1:2,000, Calbiochem, #PC205L). After washes, sections were incubated with the corresponding secondary antibody (1:1,000, AlexaFluor^®^ 488 and AlexaFluor^®^ 546 for CD68 and CGRP, respectively, Molecular Probes, USA). After PBS washes, sections were then cover-slipped with fluorescent mounting medium (Dako) and observed with Nikon Eclipse Ni-E microscope. Quantitative analyses were performed with NIS-Elements software and at least 3-6 sections per joint per animal (n=4-5 per group) were quantified using regions of interest (ROIs) on tibiotarsal joint. Results are presented as the mean intensity of signal per joint area in one ROI (2500×2500μm) for CD68 and in two ROIs (570×570μm) for CGRP.

Transverse sections (30 μm) of the brain containing the amygdala were cut on a cryotome (Microm HM450, Thermo Scientific). Free floating sections of the amygdala were stained for c-Fos immunohistochemistry as follows: after 3 washes in TBS 0.05M pH 7.6, sections were incubated for 1 h in a blocking solution (TBS 0.05M, BSA 3%, Triton 0.4%, Donkey serum 1%) and then overnight at 4°C with a rabbit primary antibody anti-Fos (1:2,000 in TBST; Santa Cruz, USA). After 3 washes in TBS 0.05M, sections were incubated for 2 h with the appropriate secondary antibody (AlexaFluor^™^ 488 goat anti-rabbit IgG; 1:1,000; Molecular Probes, USA). Sections were then washed in TBS 0.05M, mounted on gelatine coated slides and cover-slipped with Dako fluorescent mounting medium. From each animal, 3-6 sections were randomly selected for counting c-Fos positive cells in the ipsilateral side of the BLA (n=4 per group) observed with Nikon Eclipse Ni-E microscope by a blinded investigator and an average of these counts was taken.

### Electronic microscopy

Potential demyelination of the saphenous nerve was assessed at day 7, after the two knee injections of 20 nmol of LPC16:0 or vehicle using electronic microscopy as previously described^50^. Briefly, 4 animals in each group were terminally anesthetized and perfused with PAF4%/2.5% glutaraldehyde (diluted with 0.1 M sodium cacodylate buffer). A segment of the saphenous nerve was isolated proximal to the ipsilateral knee joint from vehicle and LPC16:0 groups and processed for electronic microscopy in order to evaluate the G ratio. This was calculated using the equation 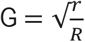 where r is the internal axonal radius, and R is the total axonal radius.

### Quantitative PCR

At day 14 and 28, 4 to 6 mice from each of the different groups (LPC16:0 10 nmol at day 14 and 28, CFA only at day 14 or vehicle at day 14 and 28) were terminally anesthetized using a mixture of Ketamine/Xylazine and ankle joints were dissected out and muscles and tendons trimmed. Samples were then flash frozen in liquid nitrogen and stored at −80⁰C until use. Frozen joints were pulverized with a biopulveriser (BioSpec, USA) followed by RNA extraction with TRIzol using Tissue Lyser II (30 Hz frequency for 2 min with 2 cycles, Qiagen, Germany). Equal amounts of RNA from each sample were reverse transcribed in 20 μl to produce cDNA using High Capacity cDNA Reverse Transcription Kit (Applied Biosystems, USA) following the manufacturer’s recommendation. SYBR Green-based detection was carried out on a real-time PCR instrument (Biorad CFX96 Touch, France) for *Il6* (Interleukin-6; F-ACCGCTATGAAGTTCCTCTC, R-GTATCCTCTGTGAAGTCTCCTC), *Tnf* (tumor necrosis factor; F-GACCCTCACACTCAGATCATCTTCT, R-CCTCCACTTGGTGGTTTGCT), *Apc5* (tartrate-resistant acid phosphatase; F-CACATAGCCCACACCGTTCTC, R-TGCCTACCTGTGTGGACATGA) and *Tnfsf11* (receptor activator of nuclear factor kappa-B ligand; F-TCTCAGATGTCTCTTTTCGTCCAC, R-CTCAGTGTCATGGAAGAGCTG). In each experiment, samples were assayed in triplicate. Data were analyzed using the threshold cycle (Ct) relative quantification method. Relative quantities (RQ) were determined using the equation: RQ = 2^−ΔΔ*ct*^. RNA 18S (F-GCCGCTAGAGGTGAA, R-CATTCTTGGCAAATG) was used as housekeeping gene.

### *In vivo* imaging

Inflammation and bone remodeling were assessed *in vivo* using an IVIS Spectrum small animal imaging system (Perkin Elmer). Each mouse received 2 nmol of MMPSense 680 (150 μl; Perkin Elmer, France) or Cat K 680 FAST^™^ (100 μl; Perkin Elmer, France) by intravenous route at day 7 and day 14, after LPC16:0 (10 nmol) or vehicle, in order to assess inflammation and bone remodeling, respectively (n=5 in each group). Six hours or 24h later for Cat K 680 or MMP680, respectively, acquisition was performed under isoflurane anesthesia (1.5%). Imaging was performed using the ex/em = 675/720 nm bandpass filters. Quantification analysis was performed with Living Image Software (Perkin Elmer) using fixed-sized rectangular ROI focused on ipsilateral or contralateral tibio-tarsal joint of the hindpaws. Results are presented as the total counts of radiance efficiency (radiance photons per second per square centimeter per steradian) per incident excitation power (microwatt per square cm).

### Cell culture and transfection

HEK293 cell line was grown in DMEM medium (Lonza™ BioWhittaker™) supplemented with 10% of heat-inactivated fetal bovine serum (Biowest^™^) and 1% of antibiotics (penicillin + streptomycin, Lonza™ BioWhittaker™). One day after plating, cells were transfected with pIRES2-rASIC1a-EGFP, pIRES2-rASIC3-EGFP or pIRES2-hASIC3a-EGFP vector using the JetPEI reagent according to the supplier’s protocol (Polyplus transfection SA, Illkirch, France). Fluorescent cells were used for patch-clamp recordings 2–4 days after transfection.

### Primary culture of DRG neurons

Lumbar dorsal root ganglia (DRG L1 to L6) were rapidly dissected out from euthanized adult male C57Bl6J (Janvier Lab, at least 8 weeks of age) and ASIC3^−/−^ mice, and placed in cold Ca-free/Mg-free HBSS medium (Corning^®^) supplemented with 10mM glucose, and 5mM HEPES (pH7.5 with NaOH). After removing the roots, DRGs were enzymatically dissociated at 37°C for 2 × 20min in the HBSS solution supplemented with calcium (CaCl_2_ 5 mM) and containing collagenase type II (Gibco, France) and dispase (Gibco, France). DRGs were then mechanically triturated and washed in a complete neurobasal A medium (NBA, Gibco^®^ Invitrogen^™^ supplemented with 2% B27, 2mM L-glutamine and 1% antibiotics: penicillin + streptomycin, Lonza^™^ BioWhittaker^™^), before being plated in 35- mm petri dishes. Neurons were then kept in culture at 37°C for one week with 1/3 of the culture medium renewed every 2 days (complete NBA supplemented with 10ng/ml NT3, 2ng/ml GDNF, 10ng/ml BDNF, 100ng/ml NGF and 100nM retinoic acid), and were used for patch clamp experiments at least 2 days after plating.

### Patch clamp experiments

The whole cell configuration of the patch clamp technique was used to record either membrane currents (voltage clamp) or membrane potentials (current clamp). The patch pipettes were made by heating and pulling borosilicate glass tubes (Hilgenberg, Germany) with a vertical P830 puller (Narishige, Japan), so that the pipette resistances were comprised between 3 and 7MΩ. Patch pipettes were filled with an intracellular solution containing either 135 mM KCl, 2.5mM ATP-Na_2_, 2.1 mM CaCl_2_, 5 mM EGTA, 2 mM MgCl_2_, 10 mM HEPES, pH7.3 with KOH for DRG neurons, or 135 mM KCl, 5mM NaCl, 5 mM EGTA, 2 mM MgCl_2_, 10 mM HEPES, pH7.3 with KOH for HEK cells. Cells were bathed into an extracellular solution made of 145mM NaCl, 5mM KCl, 2mM CaCl_2_, 2mM MgCl_2_ and 10mM HEPES for HEK cells, and this medium was supplemented with 10mM glucose for DRG neurons. pH of extracellular solutions were adjusted to different values with NMDG-Cl, including the control pH7.4 solution and the test pH6.6 solution. During patch-clamp recordings, the cell under investigation was continuously superfused with extracellular solutions using a homemade eight-outlet system digitally controlled by solenoid valves (Warner Instruments), allowing rapid changes of the immediate cell environment from control to test solutions. Electrophysiological signals generated by patch-clamped cells were amplified and low-pass filtered at 2kHz using a Axopatch 200B amplifier (Molecular Devices, UK), digitized with a 1550 A-D/D-A converter (Molecular Devices, UK), sampled at 20kHz and stored on a computer using Clampex software (V10.7, Molecular Devices). Analyses of these electrophysiological signals were then made off-line using Clampfit software (V10.7, Molecular Devices). GFP^+^ HEK cells were considered as positively transfected when the peak amplitudes of pH6.6-evoked ASIC1a or ASIC3 currents were ≥ 200pA. Non-transfected (NT) cells corresponded to GFP^−^ cells with a pH6.6-evoked current < 200pA. DRG neurons were considered as ASIC+ when they exhibited a transient pH6.6-evoked current with a peak amplitude ≥ 20pA, otherwise they were considered as ASIC-.

### *In vivo* electrophysiological recordings of spinal dorsal horn neurons

Single unit extracellular recordings of lumbar dorsal horn neurons were made using tungsten paralyn-coated electrodes (0.5MΩ impedance, WPI, Europe). The tip of a recording electrode was initially placed at the dorsal surface of the spinal cord using a micromanipulator (M2E, France) and this initial position was set as 0μm on the micromanipulator’s micrometer. The electrode was then progressively moved down into the dorsal horn until the receptive field of a spinal neuron was localized on the ipsilateral plantar hindpaw using mechanical stimulations including non-noxious brushing and noxious pinching. In this study, experiments were focused on two types of spinal neurons that were distinguished as follow: *(i)* low threshold (LT) neurons only responding to non-noxious sensory stimulations and *(ii)* high threshold (HT) neurons that dynamically respond to noxious stimulations. The depth of spinal neurons recorded in this study was 80+/−16μm (range from 10μm to 228 μm) and 202+/−21μm (range from 10μm to 571μm) for LT and HT neurons, respectively. Activities of neurons were sampled at 20 kHz, filtered (0.3-30 kHz band pass) and amplified using a DAM80 amplifier (WPI, Europe), digitized with a A/D-D/A converter (1401 data acquisition system, Cambridge Electronic Design, Cambridge, UK), and recorded on a computer using Spike 2 software (Cambridge Electronic Design, Cambridge, UK).

### Statistical analysis

Data are presented as mean ± SEM. Significant differences between the datasets are evaluated using *p*-values, which were calculated using either parametric or non-parametric tests followed by adequate post-hoc tests (see figure legends). Statistical analyses were performed using GraphPad Prism software. Association between VAS knee, VAS global, KOOS and LPC16:0 content was assessed using analysis of variance (ANOVA) using age, gender, BMI and IL-6 as covariates. The analysis was performed using R 3.6.3. Differences between sets of data and correlations were considered significant when *p*-values were less or equal to 0.05.

## Acknowledgements

We thank Drs A. Baron, S. Diochot, J. Noel, and M. Salinas for helpful discussions, V. Friend and J. Salvi-Leyral for technical support, and V. Berthieux for secretarial assistance. We also thank Tycho Tullberg, MD., PhD, CEO of Stockholm Spine Center during the data collection period for generous support and for providing research facilities at Stockholm Spine Center. The authors thank orthopaedic surgeons Ingemar Gladh and Per Gerdin for patient recruitment and taking tissue samples during surgery at Ortho Center, Stockholm. Furthermore, we thank Carola Skärvinge, research nurse at Stockholm Spine Center for excellent logistic assistance and Azar Baharpoor, Department of Physiology and Pharmacology, Karolinska Insitutet for support and laboratory assistance.

This work was supported by the Centre National de la Recherche Scientifique (CNRS), the Institut National de la Santé et de la Recherche Médicale (INSERM), the Association Française contre les Myopathies (AFM grant #19618), the Agence Nationale de la Recherche (ANR-11-LABX-0015-01 and ANR-17-CE16-0018) and the NeuroMod Institute of University Côte d’Azur (UCA). This work was also supported by the Conseil Regional Auvergne-Rhone-Alpes (project Ressourcement S3, Arth-Innov) and Feder as well as the French government IDEX-ISITE initiative 16-IDEX-0001 (CAP 20-25). We thank the multimodal imaging platform IVIA and CICS, Clermont-Ferrand, France, for *in vivo* imaging and electronic microscopy, respectively. The study has also received funding from Stockholm County Council, Swedish research Council (K2013-52X-22,199-01-3), and Eli Lilly, USA. The research leading to these results has also received funding from the European Union Seventh Framework Programme (FP7/2007–2013) under grant agreement n° 602919 and from a generous donation from Leif Lundblad and family.

## Author Contributions

Florian Jacquot, designed, performed and analyzed behavioral experiments as well as in vivo imaging; Lauriane Delay, Agathe Bayle performed and analyzed behavioral experiment; Julie Barbier performed and analyzed the electronic microscopy and immunohistochemistry experiments; Youssef Aissouni and Alexandra Jurczak did qPCR experiments; David A. Barriere helped in experimental design and critical reading of the manuscript; Fabien Marchand conceived, designed and analyzed behavioral experiments and wrote manuscript; Camilla I. Svensson helped in qPCR experiment, experimental design and critical reading of the manuscript; Anders Hugo, diagnosed patients and collected patient samples (Swedish cohort); Aisha Ahmed, performed patient data and sample collection (Swedish cohort); Eva Kosek, PI for patient data and sample collection (Sweden) and critical reading of the manuscript; Kim Kultima and Eva Freyhult helped in data analysis related to patients and critical reading of the manuscript; Spiro Khoury and Thierry Ferreira conceived, designed and performed lipidomic experiments, and analyzed data; Bonnie Labrum designed, performed and analyzed behavioral experiments; Kevin Delanoe and Ludivine Pidoux designed, performed and analyzed electrophysiological experiments; Eric Lingueglia helped in experimental design and critical reading of the manuscript; Veronique Breuil coordinated the clinical aspects of the study (French cohort); Emmanuel Deval conceived, designed, performed and analyzed experiments, and wrote the manuscript. All authors participated in the critical reading of the manuscript and give their consent for this final draft.

