## supplementary material for "Lysophosphatidyl-choline 16:0 mediates persistent joint pain through Acid-Sensing Ion Channel 3: preclinical and clinical evidences"

**Supplementary Table 1: Subject characteristics and LPC concentrations in the knee synovial fluid of the first patients' cohort.** Sexe, age and body mass index (BMI) are reported for each osteoarthritic (OA, n=35) patients. The concentrations of the different lysophosphatidyl-choline (LPC) species (LPC16:0, LPC18:2, LPC18:0, LPC20:5, LPC20:4; LPC20:3, LPC20:2, LPC 22:6) as well as the total concentrations of LPC (Sum) in the synovial fluid of each OA patient and postmortem control (n=10) are also reported in µM.

**Supplementary Table 2: Statistical analysis of LPC16:0 concentrations in the synovial fluids of OA patients (first cohort) and their pain outcomes (VAS knee, VAS global and KOOS).** Analysis of variance (ANOVA) was made using age, gender, BMI and IL-6 as covariates.

**Supplementary Table 3: Subject characteristics and LPC concentrations in the knee synovial fluid of the second patients' cohort.** Sexe, age and body mass index (BMI) and joint pathology are reported for each patient (n=50). The concentrations of the different lysophosphatidyl-choline (LPC) species (LPC16:0, LPC18:2, LPC18:0, LPC20:5, LPC20:4; LPC20:3, LPC20:2, LPC 22:6) as well as the total concentrations of LPC (Sum) in the synovial fluid of each patient are also reported in µM. Rheumatoid arthritis (RA, n=6), osteoarthritis (OA, n=18), gout (n=5), chondrocalcinosis (CCA, n=12), psoriatic arthritis (PA, n=4) and spondyloarthritis (SPA, n=5).

**Supplementary Figure 1: LPC16:0 concentrations in knee synovial fluids from patients of the second cohort.** LPC16:0 concentrations in the synovial fluid of patients from the second cohort grouped in two main joint pathologies, *i.e.,* inflammatory rheumatisms: rheumatoid arthritis (RA) + spondyloarthritis (SPA) + psoriatic arthritis (PA), and micro-cristalline arthropathies: gout + chondrocalcinosis (CCA); $^{**}p<0.01$, Unpaired t test).

**Supplementary Figure 2: LPC injections into mouse ankle produce long-lasting pain-like behaviors associated with anxiety. a,** Effect of intra-articular ankle administrations of LPC16:0 (10 nmol), complete freund adjuvant (CFA) or vehicle on mechanical allodynia of the contralateral paw of male mice. Mechanical paw threshold was assessed using the up and down method with von Frey filament on the contralateral hindpaw from D1 to D28**.** Results are expressed as 50% mechanical threshold (n=8-16 mice per group; no significant differences, two-way ANOVA followed by a Tukey's multiple comparison test). **b,** Effect of intra-articular administrations of LPC16:0 (10 nmol), CFA or vehicle on thermal hyperalgesia in male mice. Thermal withdrawal threshold was assessed using the paw immersion test at 46°C (n=8-16 mice per group; $^{****}p<0.0001$, $^{\#\#\#\#}p<0.0001$ and $^{\&\&\&\&}p<0.0001$ for a main effect on CFA and LPC16:0 curves, respectively, as compared to Veh curve; mixed-effect analysis followed by a Tukey's multiple comparison test). **c-e,** Effect of intra-articular administrations of LPC16:0 (10 nmol), CFA or vehicle in male mice on the distance travelled in the open field test (**c**), on the number of head dips in the hole board test (**d**), and on the number of buried marble in the marble burying test (**e**). The open field test lasted 5 min and the distance travelled was automatically calculated using Ethovision XT 13 (Noldus). The hole board and the marble burying tests lasted 5 and 30 min, respectively, and were analyzed by a blind experimenter (n=8 mice per group; ***p<0.001 and ****p<0.0001, one-way ANOVA followed by a Tukey's *post hoc* test)

**Supplementary Figure 3: Intra-articular LPC16:0 injections in mice are associated to increased neuronal activity within the amygdala**. **a-b,** Representative photomicrographs of c-fos expression in the amygdala at D28, after intra-articular administrations of vehicle (**a**) or LPC16:0 (10 nmol, **b**) in the ankle joint of male mice**. c**, Quantification of c-fos positive neurons in the basolateral nucleus of the amygdala (BLA). CeA: central nucleus of the amygdala; BMA: basomedial amygdala; MeA: medial amygdala (n= 4 in each group, *p<0.05, Mann-Whitney test).

**Supplementary Figure 4: ASIC3 is crucial for the development of both pain and anxiety-like behaviors induced by ankle LPC16:0 injections in male and female mice**. **a-b,** Effect of intra-articular administrations of LPC16:0 (10 nmol) or vehicle on thermal hyperalgesia in male (**a**) and female (**b**) ASIC3$^{+/+}$ and ASIC3$^{-/-}$ mice. Thermal withdrawal threshold was assessed using the paw immersion test at 46°C (n=7-8 mice per group; ***p*<0.01 and *****p*<0.0001 for ASIC3$^{+/+}$ Veh *vs.* ASIC3$^{+/+}$ LPC16:0; $^{###}p$<0.001 and $^{####}p$< 0.0001 for ASIC3$^{-/-}$ Veh *vs.* ASIC3$^{-/-}$ LPC16/0;, $^{\&\&}p$<0.01 and $^{\&\&\&\&}p$< 0.0001 for ASIC3$^{+/+}$ LPC16:0 *vs.* ASIC3$^{-/-}$ LPC16:0; Two-way ANOVA followed by Tukey's multiple comparison tests). **c-d**, Effect of intra-articular administrations of LPC16:0 (10 nmol) or vehicle on the distance travelled in the open field test in male (**c**) and female (**d**) ASIC3$^{+/+}$ and ASIC3$^{-/-}$ mice. The open field test lasted 5 min and the distance travelled was automatically calculated using Ethovision XT 13 (Noldus). Arrows indicate LPC16:0 or vehicle administrations (n=7-8 mice per group, no significant difference, one-way ANOVA followed by Tukey's post hoc tests).

**Supplementary Figure 5: LPC16:0 induces a non-inactivating ASIC3 current in a recombinant expression system. a-b** Typical ASIC current traces recorded in patch-clamp (HP -50mV) from HEK293 cells transfected with rat ASIC3 (rASIC3, **a**), rat ASIC1a (rASIC1a, **b**) or human ASIC3 (hASIC3, **c**), following extracellular acidification from pH7.4 to pH6.6. *Insets* show the effects of LPC16:0 (5µM) applied for 30 seconds at the resting pH7.4. **d**, Analysis of the current densities measured after 10, 20 and 30 second-applications of LPC16:0 (5µM) onto HEK cells either non-transfected (Ctrl GFP-, n=10) or transfected with rASIC1a (n=7), hASIC3 (n=5) or rASIC3 (n=8). Two-way ANOVA followed by a Tukey's multiple comparison test was used (**p*<0.05 and ***p*<0.01 compared to Ctrl GFP- , $^{\#}p$<0.05 and $^{###}p$<0.001 compared to ASIC1a, $^{\&}p$<0.05 compared to hASIC3. *Inset* shows the area under curve (AUC) obtained for the different transfection conditions (Ctrl GFP-, rASIC1a, hASIC3 and rASIC3). AUC were calculated using Prism software over 30 second periods from the baseline fixed at 0 (n=5-10, **p*<0.05, ***p*<0.01 and *****p*<0.0001, one-way ANOVA followed by a Tukey's multiple comparison test).

**Supplementary Figure 6: LPC knee injections generate dose-dependent, weight bearing deficit, thermal hyperalgesia and anxiety-like behaviors, without apparent neve demyelination. a,** Effect of intra-articular knee administrations of LPC16:0 (10 and 20 nmol) or vehicle on weight bearing in male mice. Results are expressed as the weight ratio between the ipsilateral and contralateral hindpaws (n=8 mice per group; $^{**}p<0.01$ and $^{****}p<0.0001$ for Veh *vs.* LPC16:0 10 nmol; $^{\#\#}p<0.01$ and $^{\#\#\#\#}p<0.0001$ for Veh *vs.* LPC16:0 20 nmol; $^{\&}p<0.05$ and $^{\&\&\&\&}p<0.0001$ for LPC16:0 10 nmol *vs.* LPC16:0 20 nmol; Two-way ANOVA followed by a Tukey's multiple comparison test). **b,** Effect of intra-articular knee administrations of LPC16:0 (10 and 20 nmol) or vehicle on thermal hyperalgesia in male mice. Thermal withdrawal threshold was assessed using the paw immersion test at 46°C (n= 8 mice per group; $^{****}p<0.0001$ and $^{\#\#\#\#}p<0.0001$ for Veh *vs.* LPC16:0 10 nmol and Veh *vs.* LPC16:0 20 nmol, respectively; Two-way ANOVA followed by a Tukey's multiple comparison test). **c-d,** Effect of intra-articular knee administrations of LPC16:0 (10 and 20 nmol) or vehicle on the number of head dips in the hole board test **(c)** and the number of buried marble in the marble burying test **(d),** in male mice. The hole board and marble burying tests lasted 5 and 30 min, respectively, and were analyzed by a blind experimenter (n=8 mice per group; $^{***}p<0.001$ and $^{****}p<0.0001$, one-way ANOVA followed by a Tukey's multiple comparison test). **e,** Effect of intra-articular knee administrations of LPC16:0 (10 and 20 nmol) or vehicle on the distance travelled in the open field test in male mice. The open field test lasted 5 min and the distance travelled was automatically calculated using Ethovision XT 13 (Noldus; n=8 mice per group; no significant differences, one-way ANOVA test followed by a Tukey's multiple comparison test). **f,** Representative electron photomicrographs of saphenous nerve sections from knee of male mice injected with vehicle (upper left) or LPC16:0 (20 nmol, upper right) at D7. G-ratios (lower panels) were calculated as a measure of myelin thickness and separated into small (left) and large (right) diameters based on internal axonal size (n=162 to 235 fibers per animal with 4 animals per group, no significant differences, Mann-Whitney tests). **g**, Contralateral paw withdrawal thresholds (PWTs) of female and male WT mice injected twice in the knees with LPC16:0 (20 nmol) or vehicle. PWTs were assessed using a dynamic plantar aesthesiometer (n=6 per group; no significant differences; three-way ANOVA test followed by a Tukey's multiple comparison test). **h**, Ipsilateral PWTs of male WT mice injected once or twice with LPC16:0 (20 nmol, n=8-14 per group, dynamic plantar aesthesiometer, $^{****}p<0.0001$ for WT LPC16:0 *vs.* WT Veh; $^{\&\&}p<0.01$ and $^{\&\&\&\&}p<0.0001$ for WT LPC16:0/Veh *vs.* WT Veh; $^{\#}p<0.05$ and $^{\#\#\#\#}p<0.0001$ for WT LPC16:0 *vs.* WT LPC16:0/Veh; Two-way ANOVA test followed by a Tukey's multiple comparison test).

| ***patient number*** | **Sexe** | **Age (year)** | **BMI** | **Joint pathology** | **LPC concentration in synovial fluid (µM)** | | | | | | | | | |
|---|---|---|---|---|---|---|---|---|---|---|---|---|---|---|
| | | | | | **LPC 16:0** | **LPC 18:2** | **LPC 18:1** | **LPC 18:0** | **LPC 20:5** | **LPC 20:4** | **LPC 20:3** | **LPC 20:2** | **LPC 22:6** | **Sum** |
| ***1*** | F | 67 | 26,8 | OA | 42,00 | 8,87 | 8,77 | 14,79 | 5,56 | 6,30 | 6,09 | 0,20 | 1,14 | 93,72 |
| ***2*** | F | 63 | | OA | 51,74 | 15,40 | 12,32 | 18,84 | 11,63 | 11,42 | 11,33 | 0,16 | 1,74 | 134,58 |
| ***3*** | F | 71 | 21,8 | OA | 46,80 | 13,36 | 11,21 | 17,87 | 9,85 | 9,73 | 10,23 | 0,22 | 1,15 | 120,43 |
| ***4*** | M | 70 | 27,0 | OA | 36,04 | 9,03 | 7,54 | 13,91 | 6,64 | 7,10 | 8,02 | 0,12 | 1,59 | 89,98 |
| ***5*** | F | 67 | 26,6 | OA | 23,60 | 4,68 | 4,86 | 8,05 | 2,85 | 3,96 | 3,96 | 0,23 | 0,58 | 52,77 |
| ***6*** | M | 67 | 31,5 | OA | 18,42 | 4,53 | 3,97 | 6,65 | 2,02 | 2,24 | 2,40 | 0,18 | 0,48 | 40,90 |
| ***7*** | F | 68 | 26,3 | OA | 33,06 | 3,83 | 4,82 | 11,62 | 1,56 | 2,60 | 3,62 | 0,16 | 0,42 | 61,68 |
| ***8*** | F | 61 | 24,7 | OA | 12,40 | 2,92 | 2,49 | 4,03 | 1,54 | 1,72 | 1,71 | 0,19 | 0,50 | 27,50 |
| ***9*** | M | 67 | 25,2 | OA | 38,10 | 11,76 | 8,95 | 10,37 | 6,69 | 5,72 | 4,60 | 0,43 | 1,11 | 87,74 |
| ***10*** | F | 73 | 25,7 | OA | 27,52 | 5,42 | 4,50 | 9,48 | 4,28 | 4,31 | 4,95 | 0,40 | 1,17 | 62,02 |
| ***11*** | F | 54 | 34,2 | OA | 67,15 | 11,95 | 15,34 | 19,31 | 7,36 | 12,06 | 9,92 | 0,29 | 1,61 | 144,99 |
| ***12*** | M | 49 | 31,1 | OA | 20,92 | 5,69 | 4,25 | 5,47 | 2,39 | 2,90 | 2,03 | 0,14 | 0,37 | 44,15 |
| ***13*** | F | 67 | 27,0 | OA | 57,21 | 7,63 | 9,94 | 18,70 | 4,93 | 8,74 | 11,24 | 0,11 | 1,16 | 119,66 |
| ***14*** | M | 70 | | OA | 43,87 | 8,50 | 8,24 | 14,12 | 5,41 | 6,92 | 6,04 | 0,24 | 0,98 | 94,32 |
| ***15*** | M | 68 | 27,3 | OA | 26,66 | 7,77 | 6,20 | 9,39 | 5,29 | 5,42 | 3,57 | 0,49 | 0,87 | 65,66 |
| ***16*** | F | 61 | 32,0 | OA | 17,13 | 4,93 | 4,27 | 5,60 | 3,05 | 2,92 | 2,55 | 0,31 | 0,53 | 41,29 |
| ***17*** | M | 73 | 28,0 | OA | 69,79 | 10,79 | 10,83 | 17,98 | 7,96 | 10,44 | 10,27 | 0,19 | 1,64 | 139,88 |
| ***18*** | M | 60 | 30,4 | OA | 25,30 | 5,38 | 6,62 | 8,07 | 3,43 | 4,74 | 3,48 | 0,20 | 0,48 | 57,69 |
| ***19*** | F | 52 | 23,2 | OA | 34,94 | 6,44 | 6,35 | 7,99 | 3,82 | 4,89 | 3,88 | 0,19 | 0,59 | 69,09 |
| ***20*** | M | 69 | 25,7 | OA | 47,57 | 8,13 | 10,88 | 13,14 | 5,27 | 8,76 | 4,68 | 0,14 | 0,83 | 99,42 |
| ***21*** | M | 60 | 27,2 | OA | 31,87 | 5,83 | 5,97 | 8,81 | 3,73 | 5,03 | 4,31 | 0,23 | 0,80 | 66,57 |
| ***22*** | F | 64 | | OA | 26,02 | 5,19 | 6,18 | 12,93 | 3,33 | 5,30 | 5,32 | 0,38 | 1,00 | 65,65 |
| ***23*** | F | 61 | 27,4 | OA | 17,74 | 3,36 | 3,78 | 7,90 | 1,93 | 2,71 | 2,95 | 0,27 | 0,47 | 41,11 |
| ***24*** | F | 62 | 27,9 | OA | 36,81 | 8,08 | 7,62 | 9,86 | 4,39 | 5,33 | 3,84 | 0,19 | 0,95 | 77,07 |
| ***25*** | M | 58 | 24,3 | OA | 33,28 | 10,83 | 9,07 | 10,86 | 4,72 | 6,01 | 3,92 | 0,26 | 0,79 | 79,75 |
| ***26*** | M | 66 | 28,4 | OA | 38,14 | 7,23 | 7,22 | 14,54 | 3,51 | 4,34 | 4,69 | 0,36 | 0,88 | 80,91 |
| ***27*** | M | 69 | 31,9 | OA | 30,58 | 4,39 | 5,91 | 10,02 | 2,38 | 4,44 | 4,17 | 0,18 | 0,86 | 62,96 |
| ***28*** | M | 56 | 29,0 | OA | 19,70 | 5,91 | 5,65 | 7,01 | 3,71 | 4,42 | 3,40 | 0,30 | 0,68 | 50,77 |
| ***29*** | M | 72 | 24,6 | OA | 30,66 | 7,14 | 5,78 | 9,38 | 4,74 | 4,76 | 4,70 | 0,41 | 0,83 | 68,42 |
| ***30*** | F | 67 | 24,8 | OA | 27,97 | 5,53 | 5,63 | 8,66 | 3,03 | 4,33 | 3,78 | 0,16 | 0,78 | 59,86 |
| ***31*** | M | 66 | 36,3 | OA | 41,35 | 7,70 | 7,31 | 11,42 | 4,89 | 5,52 | 5,08 | 0,30 | 1,98 | 85,55 |
| ***32*** | M | 73 | 22,5 | OA | 23,46 | 4,84 | 5,74 | 6,76 | 2,90 | 5,15 | 3,52 | 0,15 | 0,71 | 53,21 |
| ***33*** | M | 68 | 30,1 | OA | 31,38 | 8,31 | 5,92 | 7,78 | 5,31 | 4,43 | 3,90 | 0,16 | 0,70 | 67,88 |
| ***34*** | F | 67 | 30,1 | OA | 44,50 | 6,86 | 6,22 | 13,01 | 4,37 | 5,85 | 6,92 | 0,12 | 0,97 | 88,81 |
| ***35*** | M | 63 | 33,2 | OA | 58,52 | 9,34 | 9,63 | 13,64 | 5,73 | 7,25 | 6,61 | 0,17 | 1,23 | 112,12 |
| ***1*** | | | | PM controls | 25,74 | 5,20 | 5,03 | 7,21 | 2,64 | 3,91 | 2,33 | 0,40 | 0,77 | 53,23 |
| ***2*** | | | | PM controls | 12,94 | 1,76 | 2,58 | 5,19 | 1,02 | 1,87 | 1,33 | 0,22 | 0,37 | 27,28 |
| ***3*** | | | | PM controls | 12,31 | 2,65 | 3,28 | 6,34 | 1,60 | 2,69 | 1,45 | 0,35 | 0,59 | 31,26 |
| ***4*** | | | | PM controls | 17,66 | 11,93 | 6,08 | 5,46 | 7,30 | 5,24 | 2,82 | 0,20 | 0,71 | 57,41 |
| ***5*** | | | | PM controls | 13,71 | 2,65 | 3,85 | 3,80 | 1,97 | 4,47 | 2,15 | 0,39 | 0,65 | 33,64 |
| ***6*** | | | | PM controls | 10,55 | 2,59 | 2,54 | 4,58 | 1,29 | 1,59 | 1,07 | 0,24 | 0,32 | 24,76 |
| ***7*** | | | | PM controls | 21,58 | 4,33 | 4,54 | 5,74 | 2,05 | 2,93 | 1,81 | 0,26 | 0,54 | 43,77 |
| ***8*** | | | | PM controls | 15,68 | 5,50 | 4,13 | 10,82 | 3,12 | 3,11 | 2,08 | 0,47 | 0,93 | 45,85 |
| ***9*** | | | | PM controls | 18,83 | 3,69 | 3,49 | 6,32 | 2,23 | 2,51 | 2,44 | 0,23 | 0,89 | 40,64 |
| ***10*** | | | | PM controls | 14,00 | 3,11 | 3,67 | 10,98 | 2,56 | 3,59 | 2,68 | 0,82 | 0,81 | 42,22 |

**Supplementary Table 1**| Cohort of patients with OA (first cohort, 35 patients) and post-mortem controls (n=10).

| | *Df* | *Sum Sq* | *Mean Sq* | *F value* | *Pr(>F)* |
|---|---|---|---|---|---|
| **VASknee** | **1** | **268,6853147** | **268,6853147** | **6,681551328** | **0,016250218** |
| Gender | 1 | 4,056135654 | 4,056135654 | 0,100866244 | 0,753539278 |
| Age | 1 | 29,27231263 | 29,27231263 | 0,727931333 | 0,401989662 |
| BMI | 1 | 41,38427425 | 41,38427425 | 1,029126407 | 0,32048253 |
| log2IL6 | 1 | 76,66705017 | 76,66705017 | 1,906523367 | 0,180074114 |
| Residuals | 24 | 965,1123273 | 40,21301364 | #N/A | #N/A |
| | *Df* | *Sum Sq* | *Mean Sq* | *F value* | *Pr(>F)* |
| **VASglobal** | **1** | **220,9059633** | **220,9059633** | **5,232414512** | **0,031281663** |
| Gender | 1 | 3,462192865 | 3,462192865 | 0,082006062 | 0,777054523 |
| Age | 1 | 20,99892522 | 20,99892522 | 0,497383952 | 0,487440874 |
| BMI | 1 | 38,38032914 | 38,38032914 | 0,909082707 | 0,349863012 |
| log2IL6 | 1 | 88,18017338 | 88,18017338 | 2,088649903 | 0,161328797 |
| Residuals | 24 | 1013,249831 | 42,21874295 | #N/A | #N/A |
| | *Df* | *Sum Sq* | *Mean Sq* | *F value* | *Pr(>F)* |
| **KOOS** | **1** | **176,9162648** | **176,9162648** | **3,883437194** | **0,060404618** |
| Gender | 1 | 6,938092967 | 6,938092967 | 0,15229605 | 0,699792949 |
| Age | 1 | 14,70660843 | 14,70660843 | 0,322820461 | 0,575195495 |
| BMI | 1 | 38,56010835 | 38,56010835 | 0,846421663 | 0,36672193 |
| log2IL6 | 1 | 54,6975084 | 54,6975084 | 1,200649012 | 0,284066243 |
| Residuals | 24 | 1093,358832 | 45,55661799 | #N/A | #N/A |

**Supplementary Table 2** | Statistical analysis of LPC16:0 concentrations in the synovial fluids of OA patients (first cohort) and of their pain outcomes (VAS knee, VAS global and KOOS). Gender, age, body mass index (BMI) and synovial fluid IL-6 level were used as covariates.

| *Patient number* | Sexe | Age (year) | BMI | Joint pathology | LPC concentration in synovial fluid (µM) | | | | | | | | | |
|---|---|---|---|---|---|---|---|---|---|---|---|---|---|---|
| | | | | | LPC 16:0 | LPC 18:2 | LPC 18:1 | LPC 18:0 | LPC 20:5 | LPC 20:4 | LPC 20:3 | LPC 20:2 | LPC 22:6 | Sum |
| ***1*** | F | 26 | 30,1 | RA | 82,78 | 8,62 | 10,53 | 23,77 | 4,66 | 11,14 | 10,04 | 0,30 | 2,07 | 153,91 |
| ***2*** | F | 71 | 18,1 | RA | 39,00 | 4,97 | 6,65 | 16,06 | 3,30 | 5,36 | 7,42 | 0,33 | 1,03 | 84,12 |
| ***3*** | F | 71 | 36,0 | RA | 28,29 | 3,32 | 5,85 | 14,05 | 1,65 | 4,10 | 5,46 | 0,69 | 1,11 | 64,52 |
| ***4*** | F | 47 | 35,2 | RA | 64,02 | 13,02 | 12,30 | 31,05 | 8,44 | 9,56 | 15,76 | 0,64 | 2,02 | 156,81 |
| ***5*** | F | 75 | 26,8 | RA | 27,19 | 8,43 | 7,07 | 8,72 | 4,80 | 6,03 | 4,01 | 0,25 | 0,82 | 67,31 |
| ***6*** | F | 36 | ND | RA | 36,40 | 4,24 | 7,03 | 13,00 | 2,67 | 6,35 | 5,87 | 0,31 | 1,09 | 76,97 |
| ***7*** | F | 63 | 32,0 | OA | 46,08 | 10,77 | 9,22 | 19,68 | 7,38 | 7,52 | 10,33 | 0,44 | 1,35 | 112,78 |
| ***8*** | F | 77 | 28,3 | OA | 28,75 | 5,58 | 6,26 | 9,16 | 3,96 | 5,83 | 5,59 | 0,27 | 0,90 | 66,30 |
| ***9*** | F | 74 | 25,4 | OA | 24,24 | 6,77 | 4,59 | 9,01 | 4,64 | 4,07 | 4,87 | 0,23 | 0,61 | 59,02 |
| ***10*** | M | 73 | 24,8 | OA | 29,93 | 5,58 | 4,68 | 11,79 | 4,27 | 5,66 | 7,64 | 0,22 | 1,04 | 70,81 |
| ***11*** | M | 87 | 27,2 | OA | 77,17 | 14,50 | 19,62 | 33,14 | 9,40 | 15,73 | 15,31 | 0,44 | 2,44 | 187,75 |
| ***12*** | M | 79 | 24,7 | OA | 50,03 | 9,37 | 9,89 | 15,93 | 3,06 | 5,12 | 4,19 | 0,19 | 0,78 | 98,56 |
| ***13*** | F | 71 | ND | OA | 26,94 | 7,73 | 7,08 | 10,64 | 4,32 | 4,65 | 4,55 | 0,25 | 0,71 | 66,88 |
| ***14*** | M | 80 | 27,3 | OA | 42,03 | 9,65 | 9,03 | 15,06 | 5,90 | 7,51 | 7,86 | 0,48 | 1,38 | 98,89 |
| ***15*** | F | 82 | | OA | 55,82 | 19,32 | 14,32 | 24,93 | 16,76 | 16,00 | 17,45 | 0,47 | 2,46 | 167,54 |
| ***16*** | M | 82 | 32,3 | OA | 71,07 | 11,07 | 14,10 | 29,61 | 7,17 | 11,79 | 14,19 | 0,26 | 1,70 | 160,97 |
| ***17*** | F | 55 | 40,4 | OA | 35,54 | 8,19 | 10,68 | 14,48 | 5,44 | 8,66 | 8,13 | 0,31 | 1,65 | 93,08 |
| ***18*** | M | 70 | 29,9 | OA | 35,91 | 12,71 | 9,22 | 14,54 | 8,87 | 7,72 | 7,79 | 0,33 | 0,89 | 97,98 |
| ***19*** | F | 59 | 29,2 | OA | 36,06 | 8,77 | 7,40 | 13,52 | 4,64 | 5,27 | 5,56 | 0,38 | 1,00 | 82,61 |
| ***20*** | F | 94 | 24,5 | OA | 58,35 | 7,73 | 10,24 | 16,39 | 4,90 | 8,23 | 8,32 | 0,23 | 1,26 | 115,64 |
| ***21*** | F | 78 | 33,2 | OA | 14,36 | 5,09 | 4,05 | 5,50 | 4,02 | 4,06 | 3,57 | 0,13 | 0,64 | 41,42 |
| ***22*** | F | 70 | 29,0 | OA | 39,05 | 12,16 | 9,32 | 16,43 | 6,20 | 6,85 | 6,55 | 0,33 | 1,09 | 97,98 |
| ***23*** | F | 94 | 21,8 | OA | 25,00 | 4,49 | 4,86 | 9,74 | 2,83 | 3,75 | 4,72 | 0,23 | 0,51 | 56,14 |
| ***24*** | F | 57 | 33,9 | OA | 46,79 | 9,54 | 9,55 | 17,24 | 5,82 | 7,55 | 8,09 | 0,26 | 1,34 | 106,19 |
| ***25*** | M | 74 | 23,5 | gout | 10,59 | 1,51 | 2,60 | 3,90 | 0,93 | 2,19 | 2,10 | 0,27 | 0,49 | 24,58 |
| ***26*** | F | 63 | 32,6 | gout | 74,62 | 5,81 | 12,11 | 24,35 | 2,98 | 12,36 | 10,60 | 0,23 | 1,62 | 144,66 |
| ***27*** | M | 74 | 18,7 | gout | 20,53 | 1,62 | 3,18 | 5,49 | 0,81 | 2,94 | 2,22 | 0,30 | 0,74 | 37,82 |
| ***28*** | M | 91 | 24,3 | gout | 7,20 | 1,31 | 2,20 | 2,90 | 0,80 | 2,08 | 1,45 | 0,22 | 0,53 | 18,70 |
| ***29*** | M | 64 | 28,7 | gout | 49,48 | 4,16 | 7,59 | 14,21 | 2,43 | 6,71 | 7,21 | 0,33 | 1,27 | 93,38 |
| ***30*** | F | 81 | 23,4 | CCA | 13,88 | 1,45 | 2,97 | 6,26 | 0,86 | 2,35 | 2,60 | 0,39 | 0,50 | 31,26 |
| ***31*** | M | 83 | ND | CCA | 49,08 | 4,51 | 8,12 | 15,74 | 1,95 | 5,44 | 5,30 | 0,68 | 1,21 | 92,02 |
| ***32*** | F | 67 | ND | CCA | 25,93 | 8,53 | 5,05 | 9,52 | 5,18 | 3,78 | 4,51 | 0,24 | 0,87 | 63,60 |
| ***33*** | F | 83 | 17,8 | CCA | 53,61 | 9,02 | 12,10 | 18,99 | 5,09 | 8,78 | 7,80 | 0,44 | 1,03 | 116,86 |
| ***34*** | F | 66 | 22,6 | CCA | 46,49 | 4,72 | 8,72 | 20,89 | 2,84 | 6,95 | 10,06 | 0,29 | 1,07 | 102,04 |
| ***35*** | F | 84 | 24,7 | CCA | 5,84 | 0,81 | 1,66 | 2,27 | 0,41 | 1,09 | 0,95 | 0,23 | 0,29 | 13,55 |
| ***36*** | M | 69 | 24,6 | CCA | 15,05 | 1,97 | 2,68 | 7,07 | 1,24 | 2,54 | 4,30 | 0,37 | 0,61 | 35,81 |
| ***37*** | M | 79 | 21,0 | CCA | 30,21 | 3,46 | 5,73 | 9,41 | 2,08 | 4,84 | 5,51 | 0,40 | 0,64 | 62,28 |
| ***38*** | M | 71 | 40,2 | CCA | 20,90 | 2,39 | 5,34 | 7,53 | 1,55 | 5,25 | 4,68 | 0,29 | 0,68 | 48,61 |
| ***39*** | M | 75 | 24,8 | CCA | 54,96 | 7,73 | 11,26 | 15,24 | 5,98 | 11,12 | 10,47 | 0,44 | 1,69 | 118,88 |
| ***40*** | M | 73 | ND | CCA | 4,66 | 0,83 | 1,26 | 1,66 | 0,43 | 0,82 | 0,68 | 0,24 | 0,32 | 10,90 |
| ***41*** | F | 85 | ND | CCA | 32,75 | 4,69 | 6,30 | 9,59 | 3,29 | 5,45 | 5,59 | 0,19 | 0,82 | 68,66 |
| ***42*** | F | 50 | 20,7 | PA | 49,91 | 6,67 | 9,57 | 18,54 | 4,81 | 8,82 | 10,07 | 0,46 | 1,10 | 109,94 |
| ***43*** | F | 66 | 30,1 | PA | 59,22 | 14,56 | 13,30 | 16,07 | 11,02 | 11,60 | 9,82 | 0,18 | 1,52 | 137,29 |
| ***44*** | M | 68 | 26,1 | PA | 51,78 | 17,14 | 12,40 | 19,49 | 11,30 | 9,87 | 10,39 | 0,30 | 1,62 | 134,29 |
| ***45*** | M | 59 | 26,0 | PA | 42,83 | 4,87 | 7,19 | 16,81 | 2,66 | 5,13 | 7,63 | 0,30 | 0,92 | 88,35 |
| ***46*** | F | 55 | 34,7 | SPA | 72,94 | 6,49 | 9,31 | 24,59 | 4,04 | 7,56 | 10,83 | 0,35 | 1,59 | 137,69 |
| ***47*** | F | 41 | 20,8 | SPA | 109,87 | 9,82 | 16,43 | 44,07 | 6,96 | 15,16 | 21,20 | 0,38 | 2,34 | 226,23 |
| ***48*** | F | 33 | 28,0 | SPA | 38,73 | 6,59 | 6,66 | 17,12 | 4,50 | 6,63 | 9,08 | 0,24 | 0,97 | 90,52 |
| ***49*** | F | 48 | 22,5 | SPA | 74,26 | 13,76 | 13,14 | 32,42 | 6,99 | 8,68 | 11,51 | 0,42 | 0,98 | 162,17 |
| ***50*** | F | 57 | 26,4 | SPA | 60,79 | 6,54 | 9,34 | 21,62 | 4,10 | 10,07 | 12,85 | 0,36 | 1,66 | 127,33 |

**Supplementary Table 3** | Cohort of patients with different joint pathologies (second cohort, 50 patients, RA = rheumatoid arthritis, OA = osteoarthritis, CCA = chondrocalcinosis, PA = arhritis, SPA = spondyloathritis).

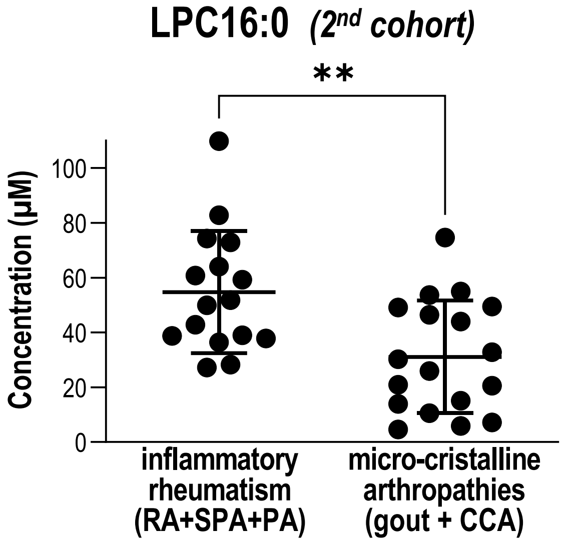

### Supplementary Figure 1

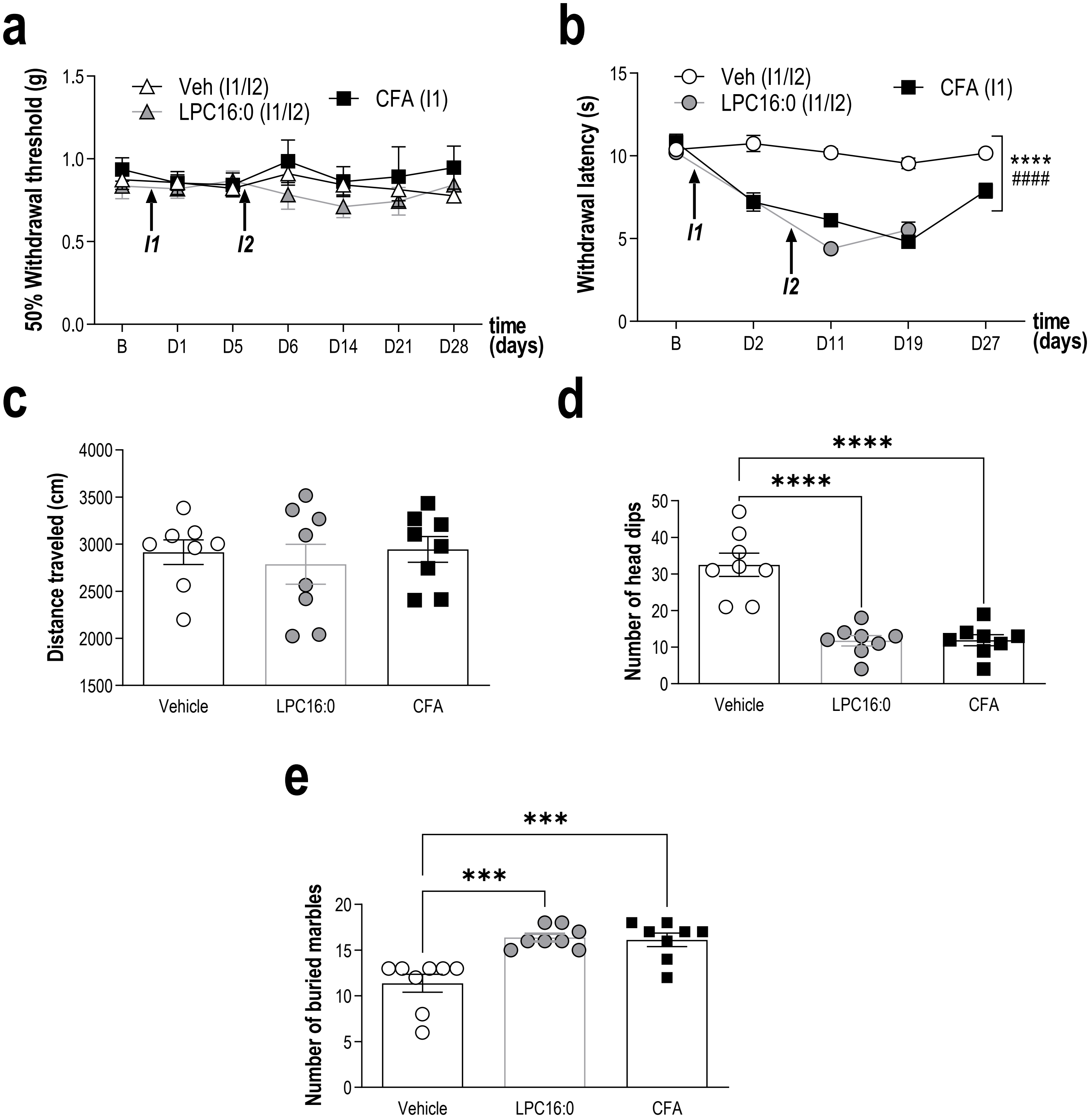

**Supplementary Figure 2**

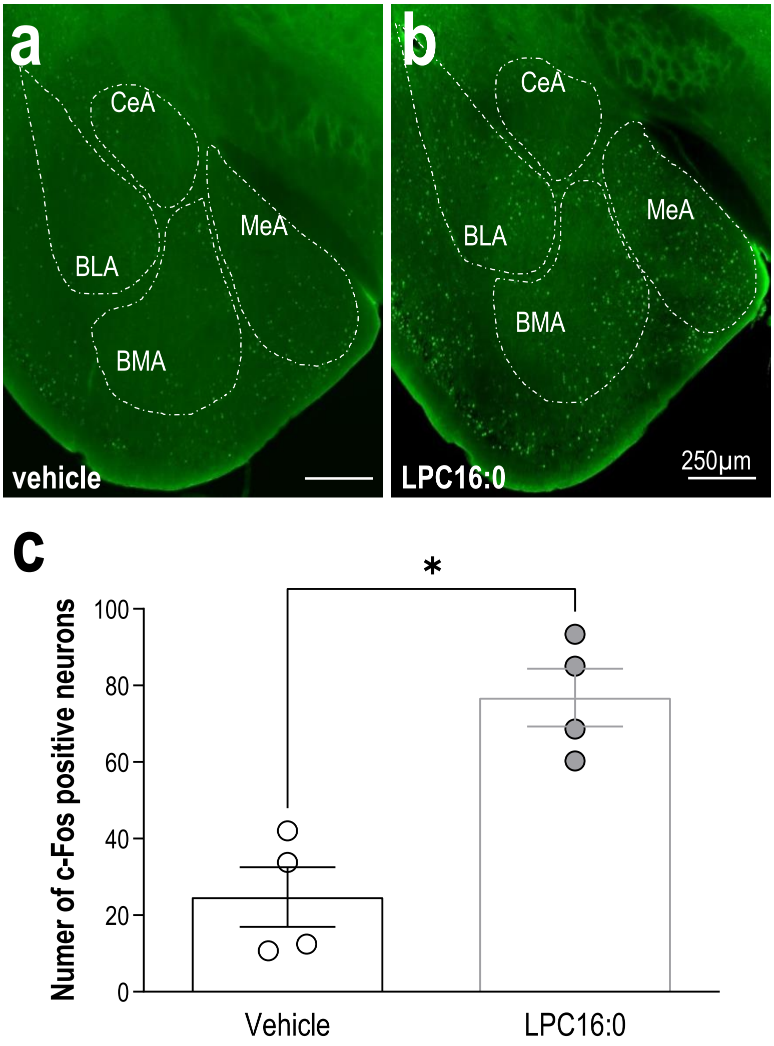

**Supplementary Figure 3**

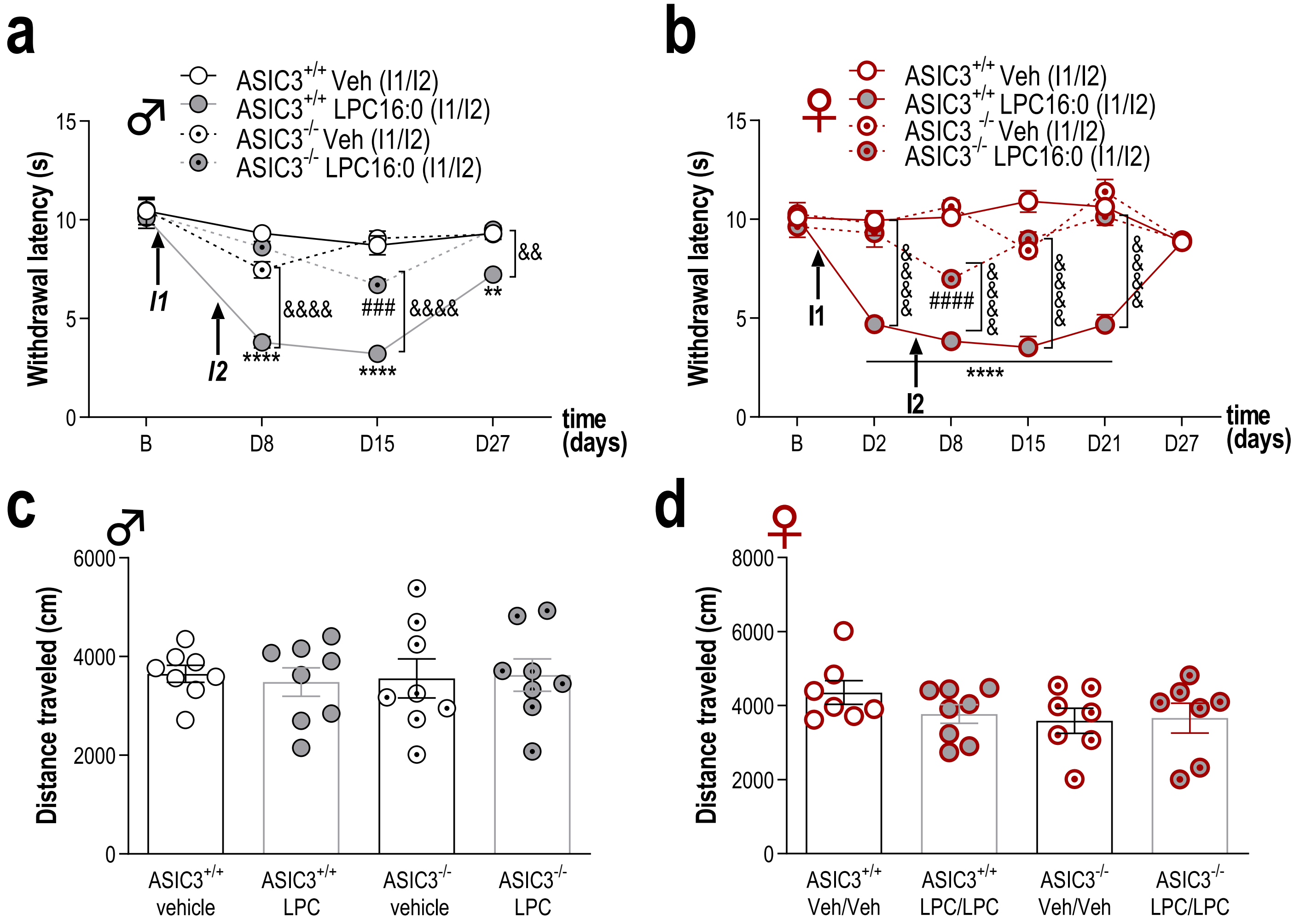

**Supplementary Figure 4**

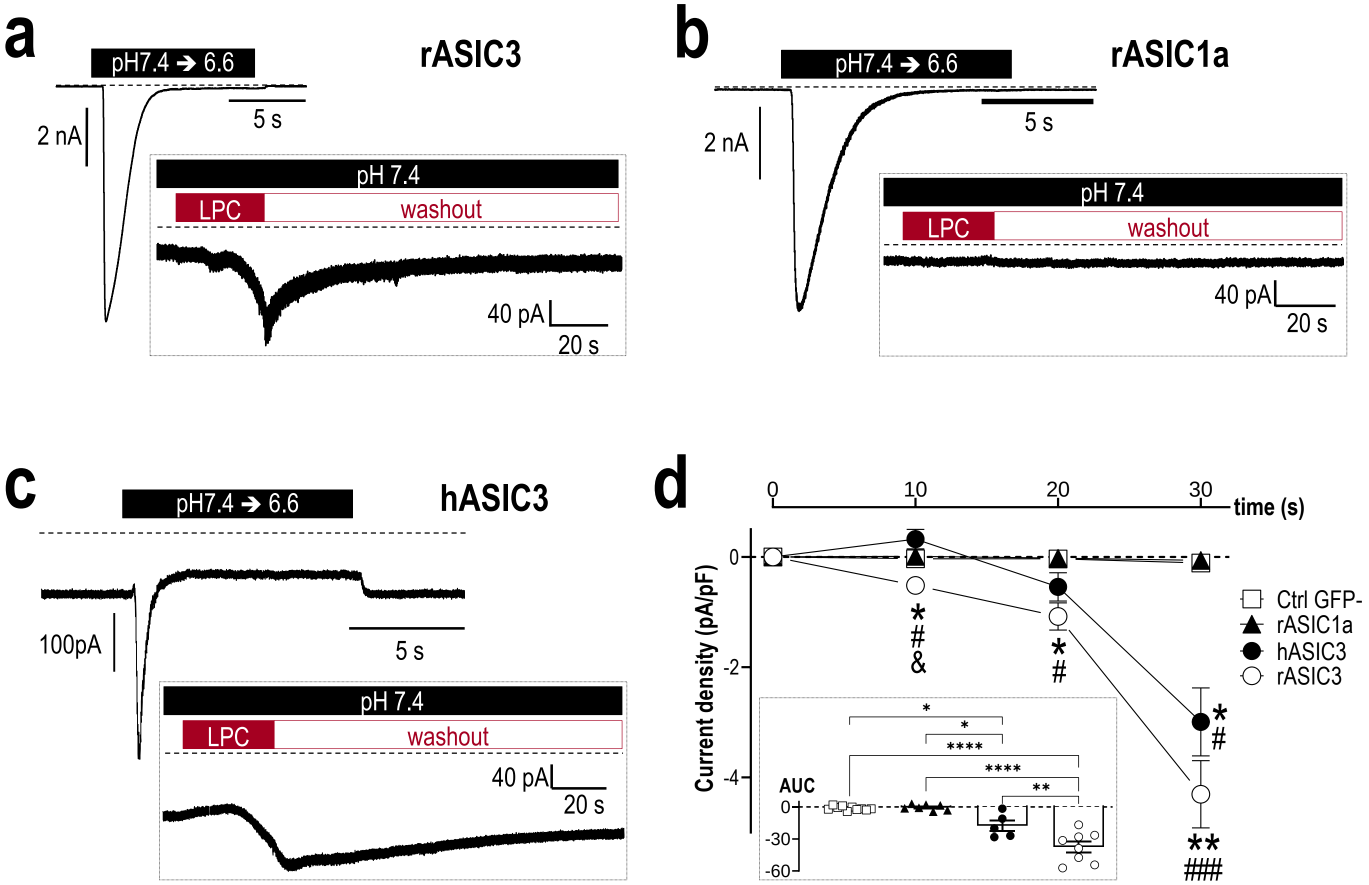

**Supplementary Figure 5**

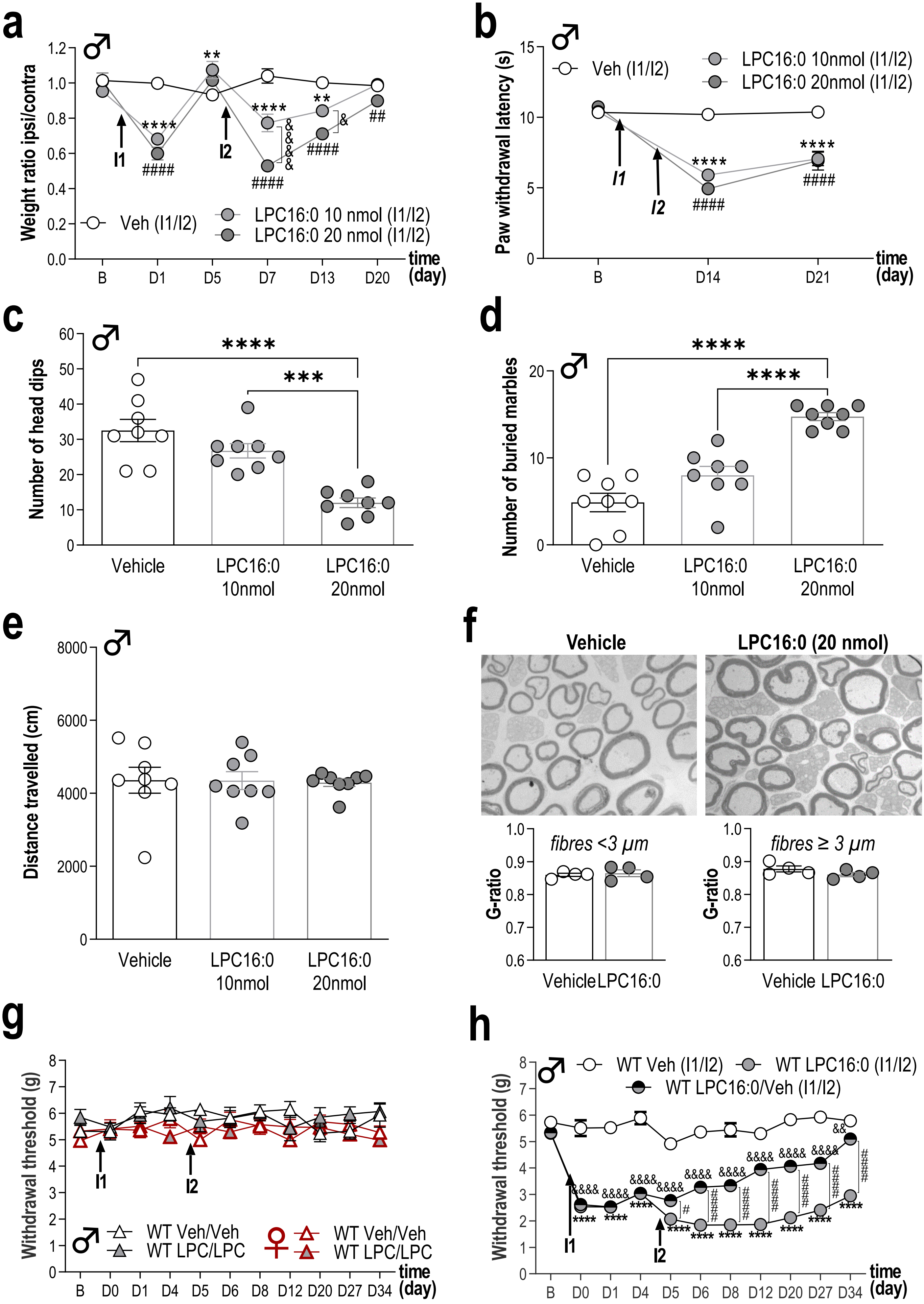

**Supplementary Figure 6**
